# Improved synthesis and application of an alkyne-functionalized isoprenoid analogue to study the prenylomes of motor neurons, astrocytes and their stem cell progenitors

**DOI:** 10.1101/2024.03.03.583211

**Authors:** Kiall F. Suazo, Vartika Mishra, Sanjay Maity, Shelby A. Auger, Katarzyna Justyna, Alex Petre, Linda Ottoboni, Jessica Ongaro, Stefania P. Corti, Francesco Lotti, Serge Przedborski, Mark D. Distefano

## Abstract

Protein prenylation is one example of a broad class of post-translational modifications where proteins are covalently linked to various hydrophobic moieties. To globally identify and monitor levels of all prenylated proteins in a cell simultaneously, our laboratory and others have developed chemical proteomic approaches that rely on the metabolic incorporation of isoprenoid analogues bearing bio-orthogonal functionality followed by enrichment and subsequent quantitative proteomic analysis. Here, several improvements in the synthesis of the alkyne-containing isoprenoid analogue C15AlkOPP are reported to improve synthetic efficiency. Next, metabolic labeling with C15AlkOPP was optimized to obtain useful levels of metabolic incorporation of the probe in several types of primary cells. Those conditions were then used to study the prenylomes of motor neurons (ES-MNs), astrocytes (ES-As), and their embryonic stem cell progenitors (ESCs), which allowed for the identification of 54 prenylated proteins from ESCs, 50 from ES-MNs and 84 from ES-As, representing all types of prenylation. Bioinformatic analysis revealed specific enriched pathways, including nervous system development, chemokine signaling, Rho GTPase signaling, and adhesion. Hierarchical clustering showed that most enriched pathways in all three cell types are related to GTPase activity and vesicular transport. In contrast, STRING analysis showed significant interactions in two populations that appear to be cell type dependent. The data provided herein demonstrates that robust incorporation of C15AlkOPP can be obtained in ES-MNs and related primary cells purified via magnetic-activated cell sorting allowing the identification and quantification of numerous prenylated proteins. These results suggest that metabolic labeling with C15AlkOPP should be an effective approach for investigating the role of prenylated proteins in primary cells in both normal cells and disease pathologies, including ALS.

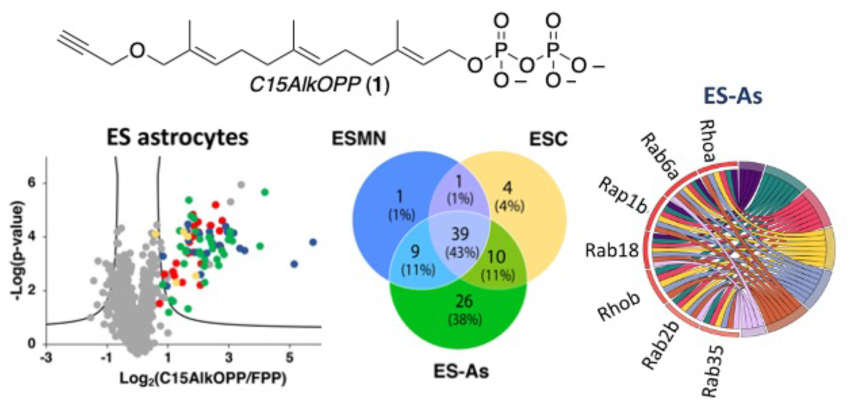

## Introduction

Protein prenylation is one example of a broad class of post-translational modifications where proteins are covalently linked to various hydrophobic moieties. Prenylated proteins incorporate either C15 (farnesyl) or C20 (geranylgeranyl) isoprenoids derived from farnesyl (FPP) or geranylgeranyl diphosphate (GGPP), respectively (Figure 1).^1^ The lipidation reaction is catalyzed by protein prenyltransferases, which transfer the isoprenoid to the thiol group of a cysteine residue located near the C-terminus of a protein substrate, resulting in the formation of a thioether bond (Figure 1). Prenylation increases the hydrophobicity of proteins, resulting in their translocation to various membranes where they participate in signal transduction pathways. However, isoprenyl groups can serve to mediate protein-protein interactions and function in other roles as well.^2^

**Figure 1.**
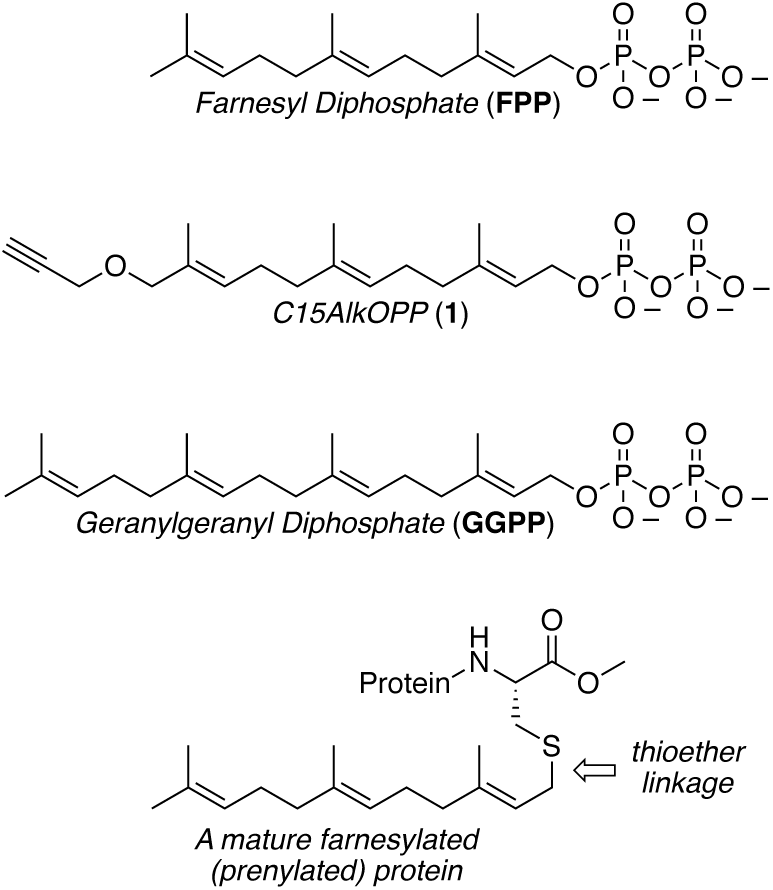
Structures of natural prenylation substrates, FPP and GGPP, the alkyne-containing probe C15AlkOPP (**1**) and the C-terminal structure of a mature prenylated protein.

Interest in prenylation initially resulted from the discovery that Ras proteins are farnesylated and that inhibition of farnesylation of oncogenic Ras variants reverses tumor formation.^3^ Those observations facilitated the development of enzyme inhibitors that subsequently moved into clinical trials;. Although none of these drug candidates have been approved for cancer therapy thus far, one of them, Tipifarnib received a Fast Track Designation from the FDA for the treatment of patients with head and neck squamous cell carcinoma in 2019.^4^ In addition, they have shown promise in other diseases. In 2020, the first farnesyltransferase inhibitor (FTI), lonafarnib, was approved for treating progeria, a premature aging disease in children, and FTIs have been developed with antifungal activity.^5, 6^ Beyond Ras-driven cancers, prenylated proteins have been implicated in the deployment of a wide variety of other diseases ranging from leukemia, Alzheimer’s disease, and amyotrophic lateral sclerosis (ALS).^7–9^

While many prenylated proteins have been identified using traditional biochemical techniques, it is likely that all of them have yet to be detected. Additionally, when prenylated proteins are implicated in a disease process, typically deduced from experiments performed with a broad spectrum of prenyltransferase inhibitors, identifying the specific involved prenylated proteins is often more complex. To address those issues and develop general methods to globally visualize, identify, and monitor all prenylated proteins simultaneously, our laboratory and others have developed chemical proteomic approaches.^10–14^ For prenylation, such methods generally rely on the metabolic incorporation of isoprenoid analogs bearing bio-orthogonal functionality. After incorporation *in cellulo*, the lysate is subjected to a bio-orthogonal biotinylation reaction, resulting in the selective labeling of all prenylated proteins. Subsequent enrichment followed by quantitative proteomic analysis allows the detection of many prenylated proteins. Work by Tate and coworkers reported the identification of 80 prenylated proteins from EA.hy926 cells using experiments performed with a combination of two different probes, one selective for farnesylation and one for geranylgeranylation.^11^ Our group reported the identification of 78 prenylated proteins in COS-7 cells using a single probe, C15AlkOPP, a substrate for both the farnesylating and geranylgeranylating enzymes.^15^ Those studies also showed that such experiments could simultaneously monitor changes in the prenylation of all observable prenylated proteins that occur upon treatment with various pharmacological agents. Recently, we reported the use of this metabolic labeling approach *in vivo*, in a mouse animal model for Alzheimer’s disease, to study the dysregulation of protein prenylation in that disease^16^ and to study the impact of knock-down of a newly identified putative deprenylating enzyme.^17^

In work reported by Li et al., it was shown that inhibitors of protein prenylation can serve as potent neurite-outgrowth-promoting agents.^9^ The authors hypothesized that prenylation may limit axonal growth in normal and pathological situations, including early-onset forms of ALS. To unravel the molecular details involved in this process, we sought to determine whether it would be possible to characterize the prenylome in motor neurons. While chemical proteomic methods have been used to profile the prenylomes of immortalized cell lines in numerous cases, there is only one example of global profiling in primary cells.^15^ Here, we first describe efforts to improve the synthesis of C15AlkOPP. Such improvements were necessary to obtain sufficient amounts of chemical probes in a single batch required to implement global profiling of prenylated proteins in a wide variety of cell cultures and animal studies. Next, the optimization of probe incorporation in metabolic labeling experiments is described, followed by an analysis of the prenylomes of embryonic stem cells (ESCs) and motor neurons and astrocytes derived from direct differentiation of ESCs. A quantitative analysis of prenylomic enrichment data obtained via metabolic labeling was then performed to examine the molecular origins of functional differences between the developmentally related cell types.

## Results

### Synthetic Optimization

The previously reported synthesis of C15AlkOPP (**1**) can be considered to be a five-step sequence (Fig. 2).^18^ While the protection (farnesol to **2**), alkylation (**3** to **5**, see Table S1 for yields) and deprotection (**5** to **6,** see Table S2 for yields) steps are relatively efficient, the yields of the oxidation (**2** to **3**) and phosphorylation (**6** to **1**) steps are poor. Several optimizations have been made to improve those transformations. In the oxidation step, allylic oxidation of the C-10 olefin leading to C-12 hydroxylation is accompanied by a competing reaction at C-6, resulting in the formation of a number of different products (see Fig. S1); the structure of the most abundant isomeric impurity (**3b**, initial oxidation of the C-6 alkene to yield a hydroxyl group at C-8) was confirmed by 2D NMR experiments (Figs. S2-S7, Table S3). Over-oxidation of the desired C-12 alcohol (**3a**) to the corresponding aldehyde (**4a**) is also problematic. These complications resulted in a low yield of 18% when performed on a 4 g scale. This reaction was studied via LC-MS, and the production of multiple isomeric products was detected (Fig. S8). Given their comparable reactivities, it was not possible to develop conditions to promote selective oxidation of the desired C-10 alkene over the C-6 olefin. However, by performing a reduction step on the crude oxidation reaction mixture using NaBH4 prior to purification, it proved possible to eliminate the carbonyl-containing impurities and simplify the subsequent chromatographic purification of the desired C-12 alcohol (**3a**). The reduced complexity of the reaction mixture was readily observable using LC-MS analysis (Fig. S8). Thus, the conversion of **2** to **3a** was accomplished on a 4 g scale in 33% yield (see Table S4 for information on four independent reactions).

**Figure 2.**
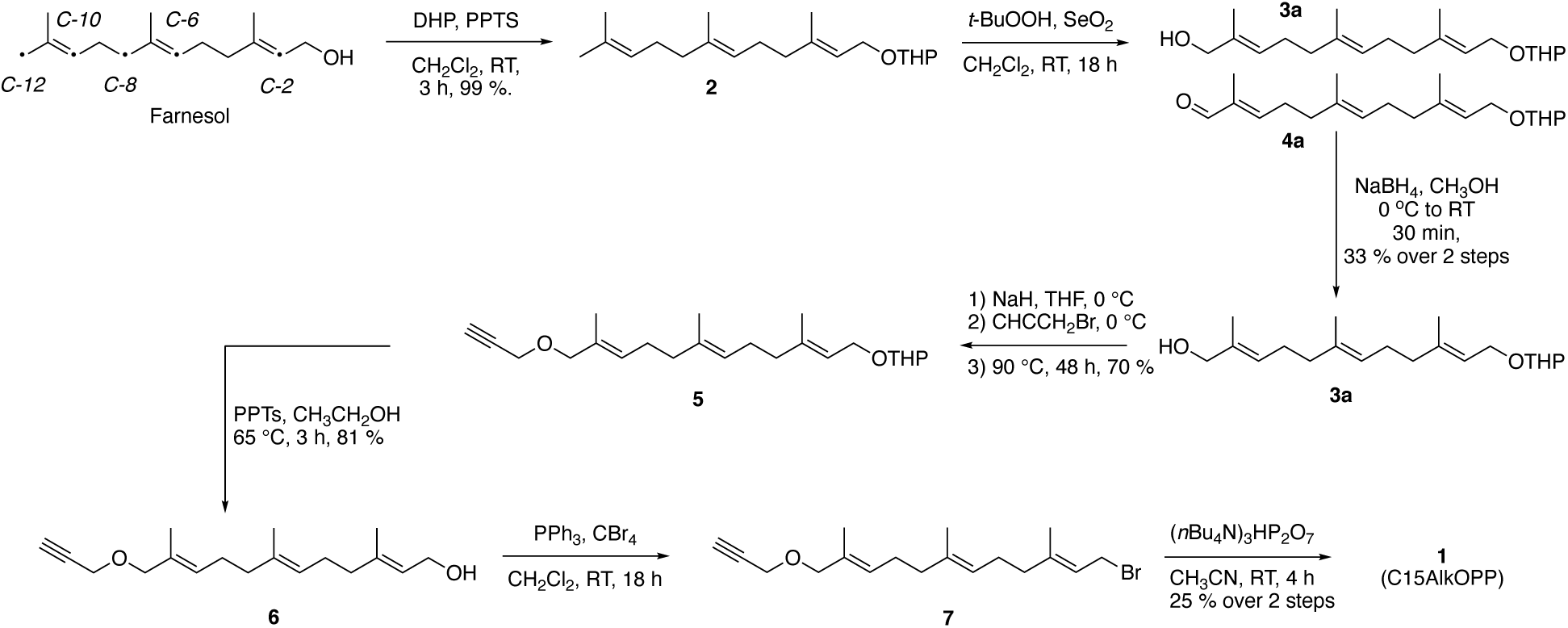
Synthetic route for the preparation of C15AlkOPP (**1**).

The conversion of alcohol **6** to diphosphate **1** is the other problematic step in this sequence. In this transformation, the starting alcohol (**6**) is first converted to the corresponding bromide (**7**), which is then used without purification, due to its potential lability, to prepare the diphosphate (**1**). Since the phosphorylating reagent is a tetra-n-butyl ammonium salt, to solubilize it in an organic solvent that will also dissolve the starting bromide, the product must be first converted to the ammonium salt via ion exchange chromatography. In our original procedure, the final diphosphate purification was accomplished via reversed-phase HPLC using a C18 column. Unfortunately, purification using that approach requires multiple injections, large amounts of solvent, and, consequently, long lyophilization times. Non-specific adsorption and decomposition on the column are also a problem in this method. Since the HPLC purification must be performed under slightly basic conditions (aqueous ammonium bicarbonate, pH 8.0) due to the instability of the product at low pH, the product is frequently contaminated with column material that has dissolved under the separation conditions. To circumvent these problems, an alternative method using cellulose chromatography was employed. Using that method, an entire reaction mixture from a ∼200 mg phosphorylation reaction could be purified using a single 2.5 x 21 cm column. With this modification, the yield of the phosphorylation reaction was 25% (see Table S5 for information on seven independent reactions). These increases in yield allowed C15AlkOPP (**1**) to be prepared in 5 % overall yield from commercially available farnesol.

### Metabolic labeling with C15AlkOPP in ES-derived motor neurons (ES-MNs)

To begin our investigations into the role of protein prenylation in motor neurons (MNs), we elected to use mouse embryonic stem cells (ESCs) because they can be efficiently differentiated into spinal motor neurons (ES-MNs) following a well-established protocol.^19,20^ The ESCs we have used express GFP under the control of the MN-specific promoter Hb9.^21^ To enrich ES-MNs, a murine ES *Hb9:GFP* reporter cell line stably transduced with a viral vector expressing also the cell surface receptor CD2 (cluster of differentiation 2) under the control of the MN-specific promoter Hb9 was used.^22^ Upon differentiation, ES-MNs were purified through a method based on magnetic-activated cell sorting (MACS). The MACS purification process gave ∼85% cell purity, following ES-MN differentiation. As a prelude to prenylomic analysis of motor neurons, their ability to undergo metabolic labeling with C15AlkOPP was first evaluated. In this approach, cells are treated with lovastatin, followed by the addition of the probe (C15AlkOPP). Statins such as lovastatin block HMG-CoA reductase, which results in the depletion of endogenous FPP production, thereby promoting the cellular machinery to incorporate the probe into the prenylated proteins in lieu of the native isoprenoid.^23^ The treated cells are then lysed and subjected to a copper-catalyzed cycloaddition of the alkyne probe with an azide-modified TAMRA-based fluorescent reagent, followed by visualization of the labeled proteins through in-gel fluorescence analysis.

We initially compared the labeling of ES-MNs to that of COS-7 and HeLa under the same protocols reported in the literature which consisted of 6 h of lovastatin treatment followed by addition of C15AlkOPP for 18 h.^15,24^ Under those conditions, it appeared that labeling in motor neurons (Fig. 3A lane 6) was significantly lower compared to COS-7 (Fig. 3A, lane 2) and Hela (Fig. 3A lane, 4), albeit similar banding patterns were observed (Fig. 3A). Therefore, we sought to explore other conditions to improve labeling in ES-MNs. While the previous methods retain the statin in the media, it appears that its presence impacts the viability of motor neurons. Accordingly, we examined alternative treatment strategies by removing lovastatin after pre-treatment for various incubation periods (Fig. 3B). Indeed, removing lovastatin from the medium allowed for longer incubation of the probe with the cells with concomitant enhancement of prenylome labeling. Maximum labeling was achieved by pre-treating the cells with 2 µM lovastatin for 24 h followed by probe incubation with 25 µM C15AlkOPP for 72 h (Fig. 3B lane 8). Hence, cell treatments in subsequent experiments were performed under those revised conditions.

**Figure 3.**
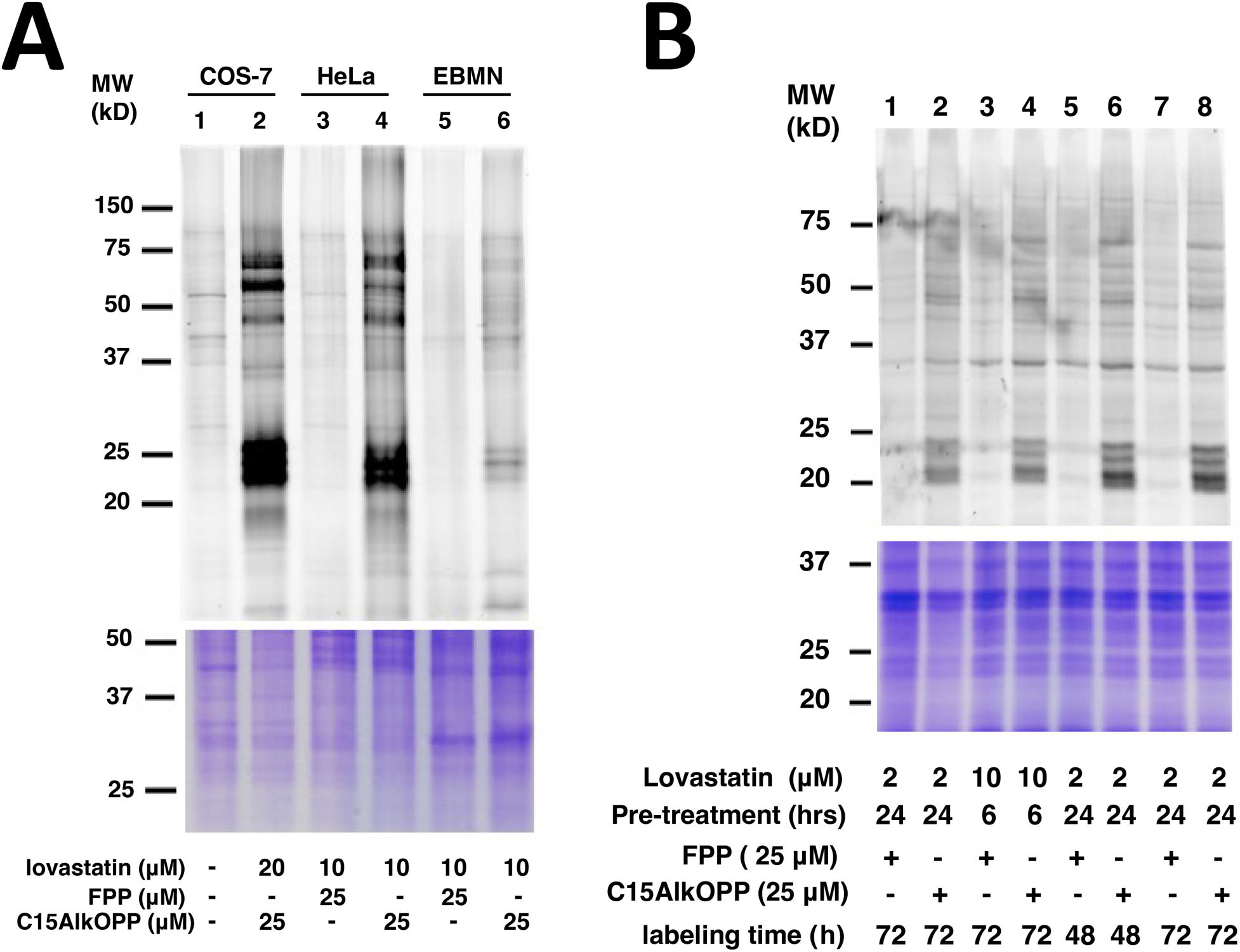
Metabolic labeling of C15AlkOPP in ES-MNs. A) Comparison of prenylome labeling in EBMNs with previously studied cell lines COS-7 and HeLa. Labeling was performed by preincubation with lovastatin at the concentrations indicated for 6 h followed by treatment with C15AlkOPP for 18 h. B) Different treatment conditions in ES-MNs to improve prenylome labeling. Labeling was performed by preincubation with lovastatin at the indicated concentrations and times (Pre-treatment) followed by treatment with 25 µM C15AlkOPP for the indicated times (probe time).

### Proteomic Analysis of ES astrocytes, ES motor neurons and ESC parent cells

ESCs can be differentiated into various cell types depending on the inducer. We were interested in how the prenylome of ES-MNs compares with that of the ESC parent cells, as well as with a related cell type, astrocytes (ES-As), which can also be differentiated from ESCs. Thus, these three cell types were subjected to metabolic labeling with C15AlkOPP under the optimized conditions described above. Samples were prepared in triplicate for the subsequent proteomic analysis of the labeled proteins (Fig. 4). Both ES-MNs (Fig. 4A) and ESC parent cells (Fig. 4B) display comparable prenylome labeling with C15AlkOPP when compared to the background FPP treatment. However, the astrocytes exhibited superior labeling, as indicated by the intense dark bands in the 25 kDa region (Fig. 4C). While both ES-MNs and astrocytes are derived from the same ESC parent cells, it is intriguing that they exhibit dramatically higher levels of metabolic labeling. This observation provided a compelling rationale for determining the identities of the labeled prenylomes.

**Figure 4.**
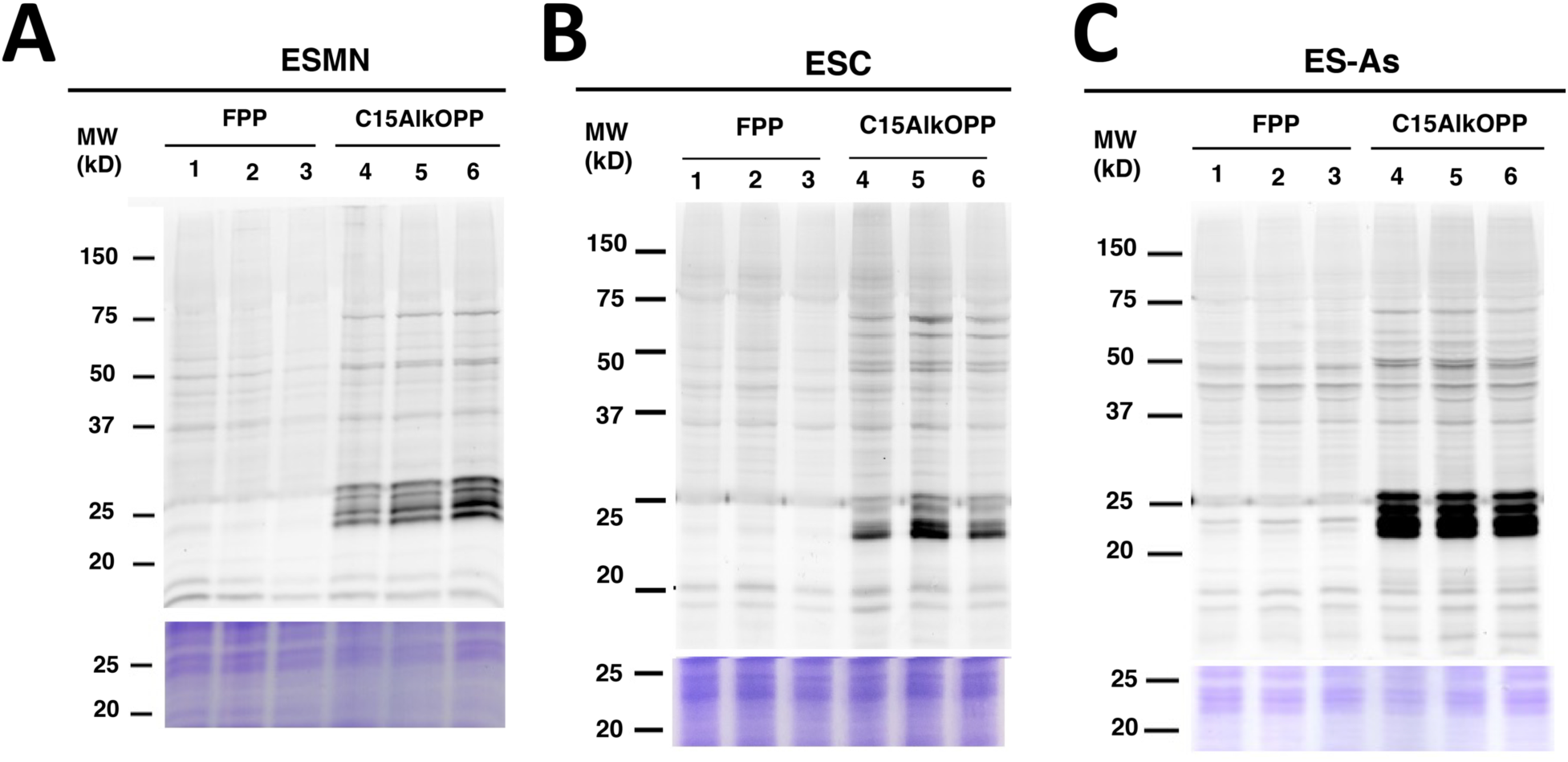
C15AlkOPP labeling in motor neurons, astrocytes and their stem cell progenitors. A) Metabolic labeling with C15AlkOPP in ES-MN cells. B) Metabolic labeling with C15AlkOPP in ESC parent cells. C) Metabolic labeling with C15AlkOPP in ES-As cells. In all cases, labeling was performed by preincubation with 2 µM lovastatin for 24 h followed by treatment with 25 µM C15AlkOPP for 72 h. In each gel, lanes 1-3 and 4-6 are from triplicate samples.

Chemical proteomic analysis was then performed in order to determine the identities of the prenylated proteins labeled by C15AlkOPP across the three cell types. Probe-treated cell lysates were tagged with biotin-azide instead of fluorophore-azide, which provides an affinity handle for subsequent enrichment. After isolating the labeled proteins, they were digested with trypsin, labeled with tandem mass tags (TMT), and subjected to mass spectrometric analysis using an MS2-based approach as previously described.^15^ From this analysis, a comparable number of prenylated proteins were enriched from ES-MNs (47 proteins) and ES parent cells (47 proteins), while almost twice the number of proteins were identified from ES-As (82 proteins) as shown in Fig. 5A; a complete list is provided in Table S6. To facilitate comparisons between datasets, protein groups that contain two or more proteins sharing an ambiguously assigned peptide sequence identified (*e. g.* isoforms) were separated into individual protein hits with the same fold change and p-values, resulting in an increased final total number of proteins identified to 50 for ES-MNs, 54 for ESCs, and 84 for ES-As. This is consistent with the metabolic labeling data shown in Fig. 4, where ES astrocytes exhibited the most intense labeling compared to the other two cell types. From the list of ungrouped identified proteins, 39 (43% of the total proteins) were common across the three cell types (Fig. 5B). Furthermore, the list of proteins identified from astrocytes included three novel proteins: Stonin-2 (CaaX box: CGVQ), Ubiquitin carboxyl-terminal hydrolase MINDY-1 or Fam63a (CaaX box: CILL), and Ubiquitin carboxyl-terminal hydrolase MINDY-2 or Fam63b (CaaX box: CVIL). While Stonin-2 is predicted to be farnesylated and both Fam63a and Fam63b proteins are predicted to be geranylgeranylated based on the web-based tool Prenylation Prediction Suite (PrePS),^25^ there is no experimental evidence for the status of their prenylation reported in the literature. This highlights the advantage of chemical proteomics in validating novel PTMs for multiple proteins in a single experiment.

**Figure 5.**
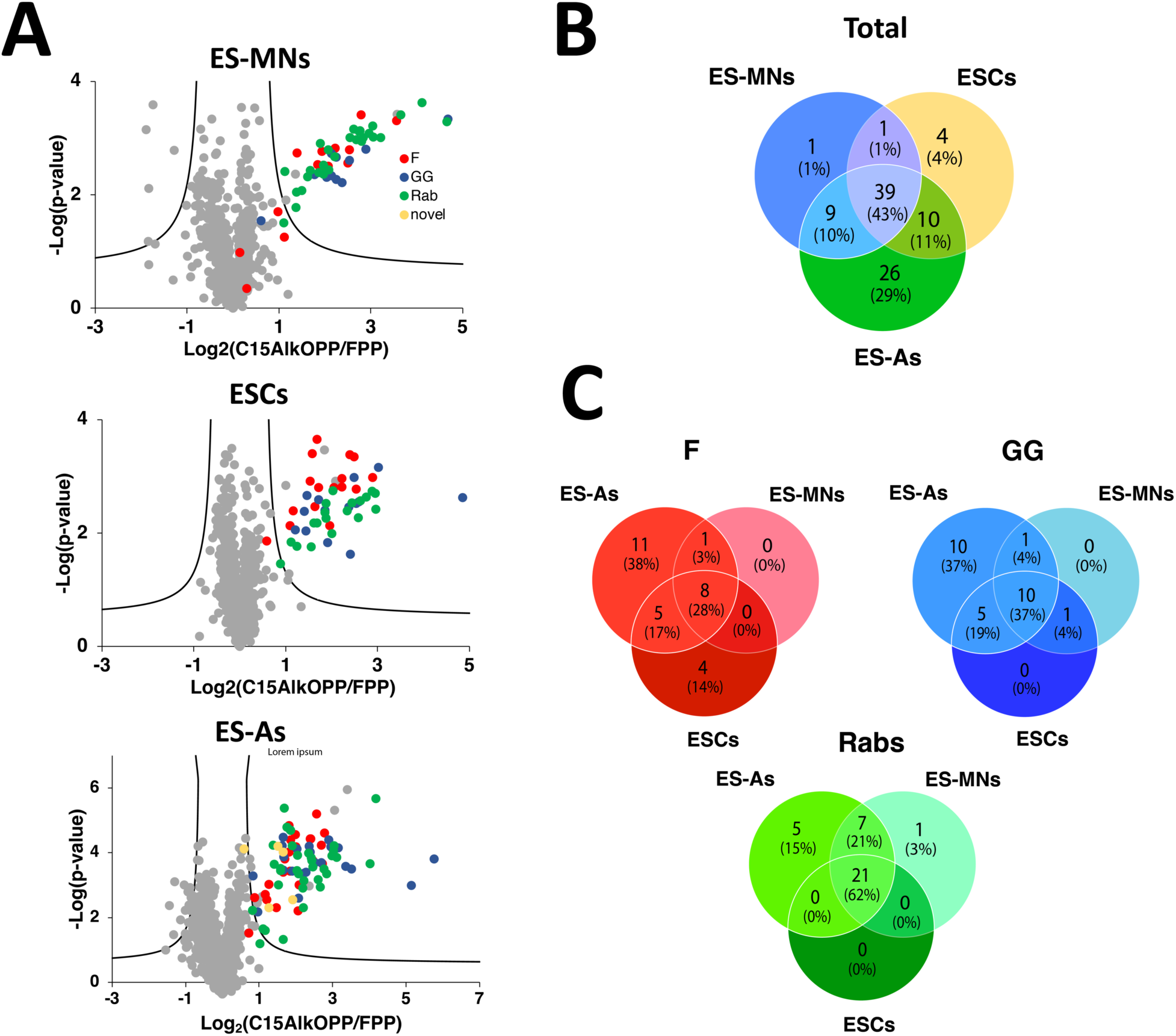
Prenylomic analysis of astrocytes, motor neurons and stem cell parent cells. A) Volcano plots (FDR = 0.01, s0 = 0.5) for the proteins identified after C15AlkOPP treatment and enrichment from ES-MNs, ESC parent cells, and ES-As. B) Comparison of the f identified proteins from the three cell types. C) Comparison of the identified proteins annotated as farnesylated (F), geranylgeranylated (GG), and Rabs. The novel proteins identified containing CaaX box motifs were assigned as farnesylated.

Interestingly, several proteins were common among all cell types, but a number of them were either shared only by two or were unique to each cell type. While comparable numbers of prenylated proteins were identified in ES-MNs and ESC parent cells, 40 were shared, and a number of unique proteins were identified from each cell type (Fig. 5B). Most of the proteins enriched in both of those cell types were also identified in ES astrocytes which provided the largest list of enriched prenylated proteins. One protein, Rab3b, was of particular interest since it was only observed in the ES-MNs; Rab3b is reported to be expressed only in neurons and is important for exocytosis.^26^ Since C15AlkOPP can be incorporated into all three classes of prenylated proteins, we investigated the distribution of C15AlkOPP labeling in each class of protein prenylation across the three cell types (Fig. 5C). It is apparent that most of the unique proteins identified from ES astrocytes were substrates of farnesyltransferase (FTase) and geranylgeranyltransferase type I (GGTase-I). Although a few Rab proteins were unique to ES astrocytes, most of them were shared among all three cell lines (62% of total Rabs identified). This may indicate that Rab proteins are more susceptible to labeling, enrichment, and detection because most of them are dually prenylated, as evidenced by the total number of detected unique proteins.

In previous work,^15,16^ we observed a correlation between transcript level and enrichment of prenylated proteins obtained via the procedure described above, suggesting that protein abundance is one of the factors that controls what prenylated proteins can be detected using this chemical proteomic workflow. Accordingly, when the fold change of shared proteins in the ES-derived motor neurons was compared to the relative transcript counts from previously reported transcriptomic work,^22^ a statistically significant positive correlation was observed (Figure S9). This same relationship between transcript levels and enrichment of prenylated proteins was previously reported in COS7 and HeLa cells.^15^ Taken together, these results emphasize the fact that the ability to detect prenylated proteins via the approach here described is, at least in part, related to their relative abundance with lower abundance proteins being more difficult to be identified.

### Comparison with previous prenylomic data

Chemical proteomic methods to label and identify prenylated proteins with the goal of defining prenylomes from various cell types have been pursued for more than a decade.^10,11,27–29^ Among these efforts, two recent studies have reported the largest lists of prenylated proteins identified through chemical tagging and enrichment to date.^11,15^ The list of grouped prenylated proteins identified from ES astrocytes reported here has the most prenylated proteins found in a single experiment for one cell line (82 proteins) inclusive of novel substrates, and is comparable to our previous analysis in COS-7 cells (78 grouped proteins) and the total proteins found in EA.hy926 human endothelial cells (88 grouped proteins) summed over multiple experiments by Storck *et al.*^11,15^ It Is important to note that the reported total proteins for COS7 and ES-As come from singular experiments, whereas the protein total for the EA.hy926 cells come from the combined results of multiple experiments, *i.e.* combined sets from probe and inhibitor dose-dependent analyses. For further analysis we compared the ungrouped prenylated proteins (84 for ES-As, 87 for COS7, and 90 for EA.hy926) enriched in each respective cell type. Subsequently, we examined the distinctions in the prenylated protein identities enriched within each specific cell type among these three different cell lines. A total of 57 proteins (50%) were enriched across the three cell types (Fig. 6A), each containing a set of unique proteins inclusive of the novel proteins identified at the time they were reported. While more unique proteins that bear CaaX boxes were seen in EA.hy926 cells, many of those proteins appeared to be unresponsive to competition assays performed in that study and were deemed unprenylatable by PrePS, including ANXA1 and RPL12, suggesting that they may not be *bona fide* prenylation substrates and appear in the prenylomic analysis due to artifactual reasons. Hence, enriching novel prenylated proteins from a chemical proteomic approach is a good step to initially interrogate the prenylation status of a protein although further validation studies are often required.

**Figure 6.**
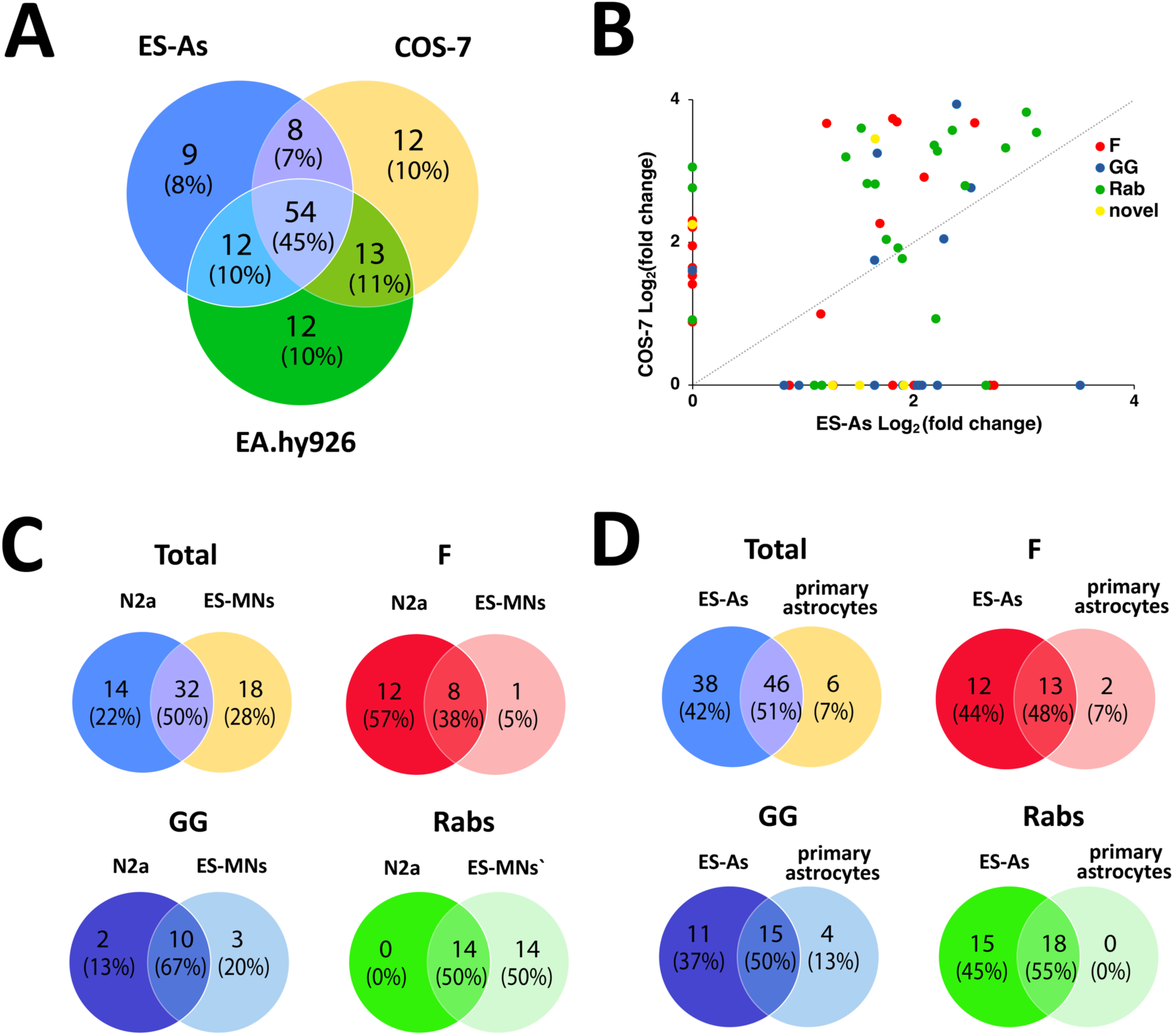
Comparison of prenylomic data obtained here with that observed in previous studies with COS-7 and E.hy296 cells. A) Comparison of the number of prenylated proteins enriched in ES-As (this work) compared with previous studies in COS-7 and EA.hy926 cells. B) Comparison of the fold change obtained for prenylated protein enrichment in ES-As (this work) and COS-7 (previous work). C) Comparison of the number of prenylated proteins and their prenylation type obtained in ES-MNs (this work) compared with previous results with N2a cells. D) Comparison of the number of prenylated proteins and their prenylation type obtained in ES-As (this work) compared with previous results with primary astrocytes. Data for COS-7, N2A and primary astrocytes is from Suazo *et al*.^15^ and data for EA.hy926 cells is from Storck *et al.*^11^

We then investigated the extent of correlation in terms of prenylation fold enrichment with individual protein levels identified between ES astrocytes and COS-7, which provided comparable numbers of total prenylated proteins. The fold enrichment values obtained for individual proteins from the two experiments were plotted to compare their extent of enrichment in each cell type (Fig. 6B). It appears that the extent of enrichment in COS-7 cells for most prenylated proteins is greater than in ES astrocytes. While both cell types displayed strong labeling in the in-gel fluorescence analyses (Fig. 3A and 4C), it is important to note that intense bands may not directly correspond to higher labeling of individual proteins, as many prenylated proteins co-migrate in one region of the gel around 25 kDa.

The above comparison of various cell types highlights the fact that substantial differences in protein labeling and enrichment are observed across different cell lines. Those results probably convolute a variety of variables, including probe uptake and prenyltransferase expression, as well as functional cell differences. Thus, we next compared prenylomes identified from more similar cell lines. The prenylomes of the neuronal cell line N2a and primary astrocytes derived from mouse brain were previously characterized.^15^ Comparison of the prenylomes determined using C15AlkOPP between ES-MNs and N2a provided distinct sets of prenylated proteins with ∼50% similarity (Fig. 6A). Although the total number of prenylated proteins identified in each cell type was comparable, more farnesylated proteins were enriched in the neuronal cell line while more Rabs were identified in motor neurons (Fig. 6C). Similarly, the stem cell-derived ES astrocytes share ∼50% with the mouse brain primary astrocytes with almost all proteins from the latter being identified in the former (Fig. 6D).

### Bioinformatic analysis of ESCs, ES-astrocytes (ES-As) and ES-motor neurons (ES-MNs)

A quantitative bioinformatic analysis was undertaken to gain additional insight into the function of the prenylated proteins identified in this study. For that, the log-fold change-ranked list of the significantly prenylated proteins (p <0.05) in each cell type was used to identify enriched biological pathways using three databases, including KEGG, Reactome, and Wikipathways by pre-ranked Over-representation analysis (ORA). Of note, significant enriched pathways (p<0.10) were detected only for ES astrocytes (Fig. 7). Using the ungrouped data set of 39 proteins common to all three cell types (ESCs, ES-MNs and ES-As), they were found to be related to nervous system development, chemokine signaling, Rho GTPase signaling, and adhesion. Next, we tested whether there are some proteins among the 39 proteins that are specifically more prenylated in one cell type over the others. Hierarchical clustering in a heatmap of the log-fold change values across cell types allowed the identification of a set of highly prenylated proteins for each specific cell type (Fig. 7A). The Over representation analysis (ORA) in Gprofiler was performed for cell type specific hyper-prenylated proteins based on heatmap clustering. The most enriched pathways in all the three cell types are related to GTPase activity and vesicular transport (Fig. 7B, Table S7). Furthermore, STRING analysis was used to highlight documented interactions among the proteins in each signature (confidence=0.07). When the ungrouped set of proteins was considered, significant interactions were found in two populations that appear to be cell type dependent (Figure 7C and Table S7).

**Figure 7.**
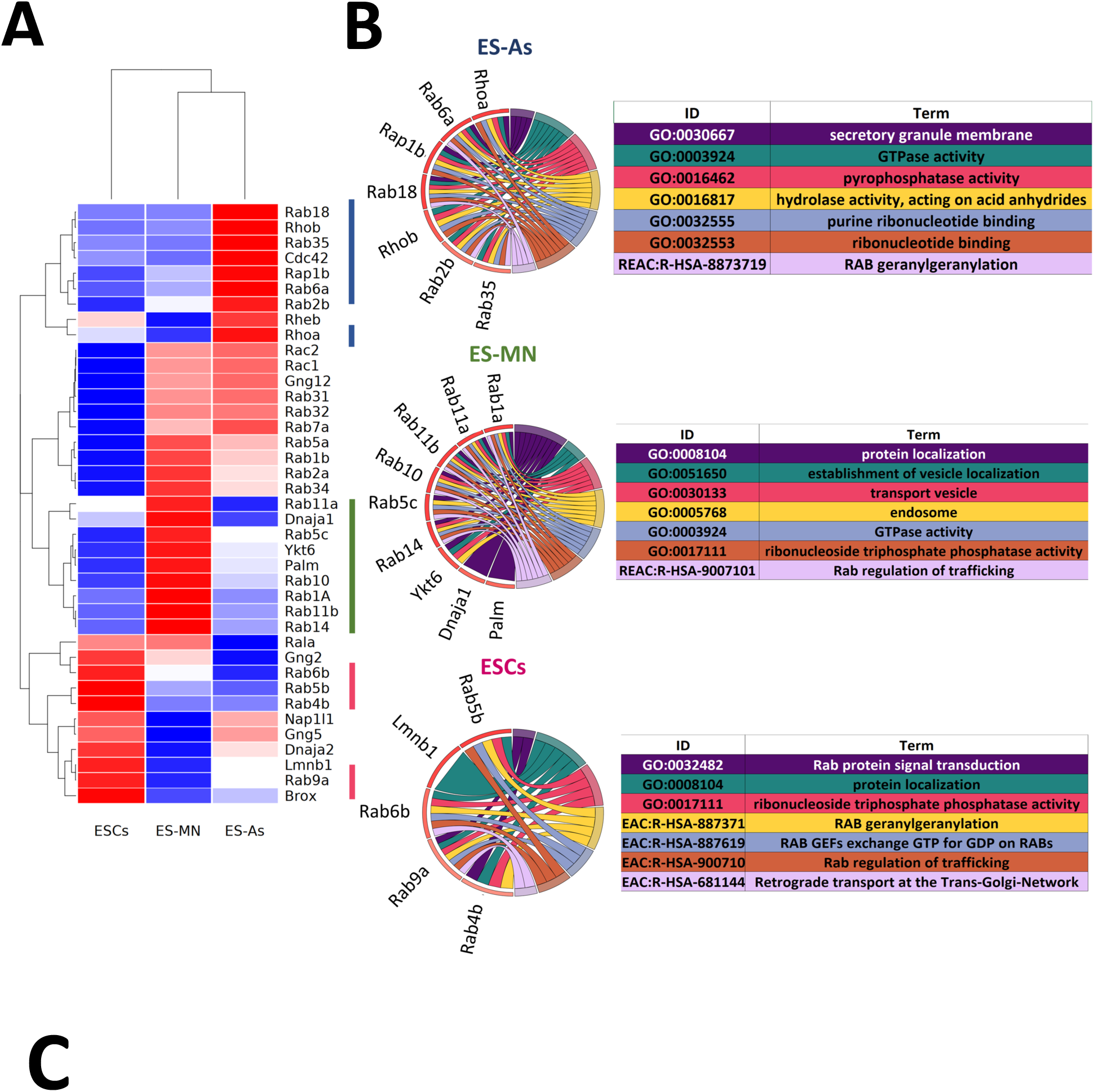
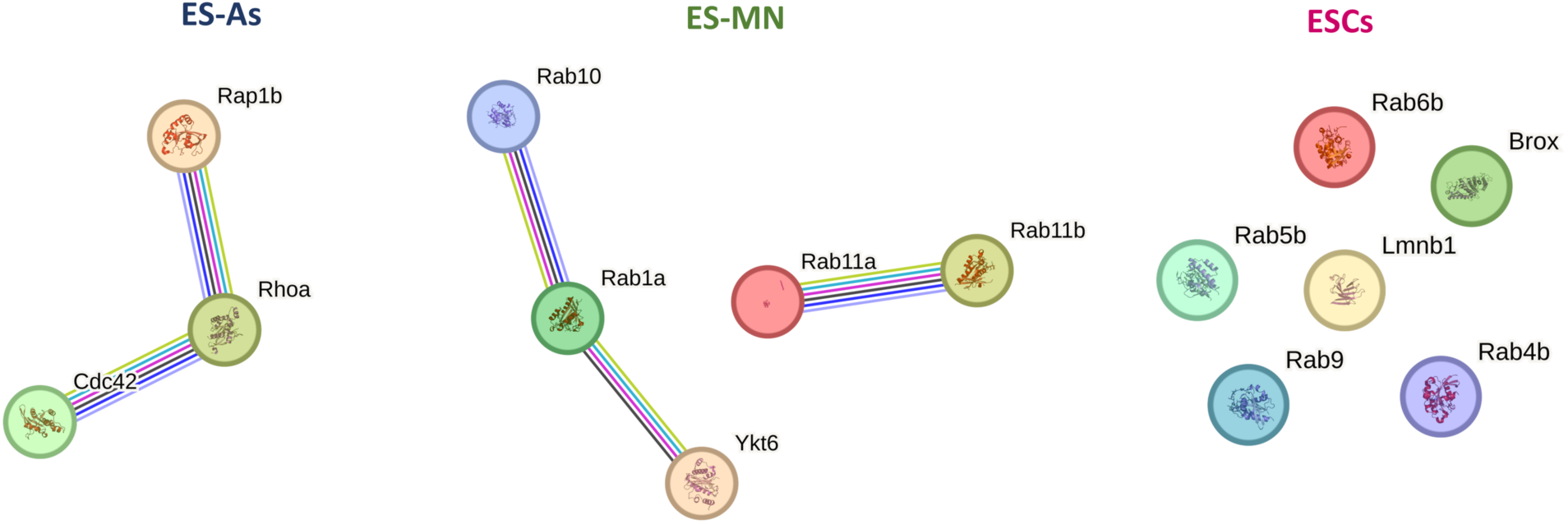
Bioinformatic analysis of ESCs, astrocytes (ES-As) and motor neurons (ES-MN). A) Heatmap of scaled prenylated proteins. B) Cord diagrams for the top 5 enriched pathways, sorted for padj, and resulting from Over Representation Analysis (ORA) performed with gProfiler R package, on the hyper-prenylated proteins for each population as indicated by the color codes bars on the right of the heatmap. Gene color is function of scaled fold change. C) STRING protein connections among selected hyperprenylated proteins.

Through the above bioinformatic analysis of the proteins found common in the astrocytes, motor neurons, and ES cells (Fig. 7), it was discovered that there were significant differences in the prenylation of specific proteins found in those cells. Interestingly, the data showed that prenylated proteins with documented roles associated either with the biological function of either astrocytes or motor neurons were found to be enriched in their respective cell types. Such an example is the increased presence of the small GTPase Cdc42 in astrocytes (Fig. 7A). In migrating astrocytes, Cdc42 is known to control cell polarity and importantly is a key regulator of vesicle trafficking.^30^ Interestingly, exploring the localization and role of the two isoforms of Cdc42 (Cdc42b or Cdc42u) showed that when Cdc42 was knocked down in cells, reconstitution with a non-prenylatable mutant of either isoform resulted in no rescue of function.^30^ This suggests that in mature astrocytes, the prenylated form of Cdc42 is critical for cellular function. Another noteworthy protein is Ytk6 which is an established prenylated protein that was found to be enriched in the motor neuron cells (Fig. 7A). As a SNARE protein, Ytk6 is crucial for synaptic vesicle exocytosis in neurons.^31^ Interestingly, unlike other related SNARE proteins which have a transmembrane domain for anchoring in the lipid membrane, Ytk6 instead has a prenylation recognition site for lipid anchoring, providing a molecular explanation for prenylation in Ytk6 given its function.^32^ Many of the other proteins that manifested cell-specific differences in their levels (Fig. 7A) are highly related small GTPases, but they also have distinct roles unique to specific cell types. When examining the Circos plots (Fig. 7B), it was noteworthy that in the motor neurons Rab11b and Rab5c were prenylated more than in the other cells in this study. Those Rab proteins mediate cellular recycling and N-Cadherin trafficking, and hence regulate neuronal migration and dendrite formation.^33^ While there are obvious reasons why some of the prenylated proteins from this study were enriched to higher levels in motor neurons and astrocytes given their documented roles in cell specific functions, differences in the ES cells are harder to explain given that less information is available. In those cells, the proteins found to be more highly enriched included Rab4b, Rab6b, Rab5b and Rab9a. What is known is that some of these proteins are essential for cellular function mainly endosome modulation.^34^ and when knocked out, result in detrimental effects.^35^ It may also be that these small GTPases are crucial housekeeping and general maintenance regulators that become less critical as the cell differentiates. Taken together, this bioinformatic analysis suggests that beyond simply identifying proteins that are present in different cell types, quantitative analysis of prenylomic data obtained via metabolic labeling and enrichment can provide additional insights into the molecular origins of functional differences between developmentally related cell types. While potentially quite powerful, there has been limited use of metabolic labeling coupled with proteomic analysis to explore changes in post-translational modification that occur during development. Thus far, most of those studies have focused on changes in protein glycosylation that occur during differentiation of various stem cells using glycans functionalized with bio-orthogonal reporters.^36–38^ To our knowledge, this is the first example of the use of metabolic labeling in concert with bioinformatic analysis to study changes in post-translational lipid modifications that occur in cellular differentiation.

## Conclusions

In this study, the synthesis of the C15AlkOPP probe was improved to facilitate chemical proteomics experiments. This goal was accomplished by optimizing the oxidation and phosphorylation steps giving an overall yield of 5% over six synthetic transformations. The material obtained from that optimization was used for metabolic labeling experiments with ESCs and motor neurons and astrocytes derived from direct differentiation of ESCs. A series of experiments with those cells showed that a 24-hour preincubation with statin followed by 72-hour treatment with probe gave optimal labeling and minimized toxicity. Chemical proteomic analysis of the proteins enriched after metabolic labeling with C15AlkOPP allowed for the identification of 5 prenylated proteins from ESCs, 50 from ES-MNs, and 84 from ES-As. That last number is the largest number of prenylated proteins identified in a single experiment. Consistent with previous observations, a correlation was observed between protein enrichment values obtained by quantitative mass spectrometry and transcript abundances obtained from transcriptomic experiments. Bioinformatic analysis revealed specific enriched pathways, including nervous system development, chemokine signaling, Rho GTPase signaling, and adhesion. Hierarchical clustering showed that most enriched pathways in all three cell types are related to GTPase activity and vesicular transport, while STRING analysis showed significant interactions in the differentiated populations. Collectively, the data provided herein demonstrates that robust incorporation of C15AlkOPP can also be obtained in ES-MNs and related cells, allowing the identification and quantification of numerous prenylated proteins. These results indicate that metabolic labeling with C15AlkOPP should be an effective approach for investigating the role of prenylated proteins in neuronal growth in both normal cells and disease pathologies, including ALS. Finally, from a global perspective, it should be noted that the successful use of metabolic labeling and proteomic analysis in specific cell types obtained via magnetic activated cell sorting suggests that this approach could have widespread utility in a plethora of other cell types.

## Materials and Methods

### General

Solvents and reagents used were purchased from Sigma-Aldrich (St. Louis, MO) and were used without further purification. TAMRA-N3 and Biotin-N3 were purchased from BroadPharm. ^1^H NMR data of synthetic compounds were recorded at 500 MHz on a Bruker Avance III HD Instrument at 25 °C. Mass spectra of organic compounds were acquired using either a Bruker BioTOF II ESI/TOF-MS, or an Applied Biosystems-Sciex 5800 MALDI-TOF instrument. TAMRA fluorescence analysis of gels was performed using a Typhoon FLA 9500 (GE Healthcare). Gel images were processed in ImageJ.

### 2-(((2E,6E)-3,7,11-trimethyldodeca-2,6,10-trien-1-yl)oxy)tetrahydro-2H-pyran (2)

To a stirred solution of farnesol (3.01 g, 13.5 mmol, 1.0 equiv) in dry CH2Cl2 (25 mL), DHP (2,3-dihydropyran, 1.72 g, 20.3 mmol, 1.5 equiv) was added at RT. Next, PPTs (351 mg, 1.35 mmol, 0.1 equiv) was added to the reaction mixture and the resulting solution was stirred overnight at RT. After 18 h the reaction was quenched with sat. aq. NaHCO3 (15 mL) and extracted with CH2Cl2 (3 x 15 mL). The combined organic layers were concentrated and the crude product (4.6 g, 99%) was directly used for the next step without purification. The average reaction yield when performed 3 times was 99%. ^1^H NMR (400 MHz, CDCl3) δ 5.41 – 5.30 (m, 1H), 5.16 – 4.99 (m, 2H), 4.61 (dd, *J* = 4.3, 2.9 Hz, 1H), 4.21 (dd, *J* = 11.9, 6.4 Hz, 1H), 4.01 (dd, *J* = 11.9, 7.4 Hz, 1H), 3.87 (ddd, *J* = 11.2, 7.6, 3.3 Hz, 1H), 3.55 – 3.38 (m, 1H), 2.09 (t, *J* = 7.6 Hz, 2H), 2.03 (dd, *J* = 8.4, 6.1 Hz, 4H), 1.99 – 1.91 (m, 2H), 1.87 – 1.76 (m, 1H), 1.69 (s, 1H), 1.68 – 1.63 (m, 6H), 1.60 – 1.55 (m, 6H), 1.54 – 1.47 (m, 4H).

### (2E,6E,10E)-2,6,10-trimethyl-12-((tetrahydro-2H-pyran-2-yl)oxy)dodeca-2,6,10-trien-1-ol (3a)

To a stirred solution of THP-protected farnesol **2** (4.01 g, 13.1 mmol, 1.0 equiv) in dry CH2Cl2 (25 mL), SeO2 (131 mg, 1.30 mmol, 0.1 equiv) and salicylic acid (179 mg, 1.30 mmol, 0.1 equiv) was added at RT. Next, *^t^*BuOOH (6.0 mL, 39 mmol, 3.0 equiv) was added dropwise to the reaction mixture at RT over 10-12 min and the resulting solution was stirred overnight at RT. After 24 h, the reaction was quenched with sat. aq. NaHCO3 (15 mL) and extracted with CH2Cl2 (3 x 15 mL). The combined organic layers were concentrated to dryness *in vacuo* and then dissolved in EtOH at 0 °C (ice bath). To the reaction mixture. 0.2 equiv (100 mg, 2.6 mmol) of NaBH4 was added at 0 °C. After 15 min, the reaction was warmed to RT and stirred for another 15 min. After removal of the solvent *in vacuo*, the resulting crude product was purified by column chromatography on silica gel (EtOAc:Hexanes, gradient of 5:95 to 20:80, v/v) to afford 1.4 g (33% yield) of the desired allylic alcohol as a viscous yellow oil. The average reaction yield when performed 4 times was 33%. That data is summarized in Table S1. ^1^H NMR (400 MHz, CDCl3) δ 5.45 – 5.28 (m, 2H), 5.11 (t, *J* = 6.1 Hz, 1H), 4.62 (dd, *J* = 4.6, 2.5 Hz, 1H), 4.28 – 4.18 (m, 1H), 4.07 – 4.00 (m, 1H), 3.98 (s, 2H), 3.93 – 3.82 (m, 1H), 3.57 – 3.36 (m, 1H), 2.21 – 2.08 (m, 4H), 2.07 – 1.97 (m, 4H), 1.90 – 1.76 (m, 1H), 1.73 (dd, *J* = 9.9, 6.1 Hz, 1H), 1.67 (s, 3H), 1.65 (s, 3H), 1.60 (s, 3H), 1.55 (d, *J* = 11.2 Hz, 4H). (SM-1003).

### 2-(((2E,6E,10E)-3,7,11-trimethyl-12-(prop-2-yn-1-yloxy)dodeca-2,6,10-trien-1-yl)oxy)tetrahydro-2H-pyran (5)

To a suspension of NaH (429 mg, 10.7 mmol, 2.5 equiv, 60% in oil) in dry THF (12 mL), a solution of allylic alcohol **3a** (1.38 g, 4.28 mmol, 1.0 equiv) in THF (3 mL) was added dropwise under N2 atmosphere at 0 °C (over 10 min). After 15 min at 0 °C, the temperature was raised to RT and the reaction mixture was stirred for another 15 min. Then, the reaction mixture was cooled to 0 °C and a solution of propargyl bromide in toluene (2.9 mL, 26 mmol, 6.0 equiv) was added dropwise under a N2 atmosphere over 5 min. Next, the reaction was warmed up to RT and then heated to 90 °C for 2 days. The reaction was quenched with cold water (15 mL) and extracted with EtOAc (3 x 10 mL). The combined organic layers were concentrated *in vacuo* and purified by column chromatography (silica gel, EtOAc:Hexanes, gradient of 0:100 to 5:95, v/v) to afford 1.01 g (65% yield) of the desired product isolated as a yellow oil. The average reaction yield when performed 5 times was 70%. That data is summarized in Table S2. ^1^H NMR (500 MHz, CDCl3) δ 5.42 (t, *J* = 7.1 Hz, 1H), 5.36 (t, *J* = 6.8 Hz, 1H), 5.11 (t, *J* = 6.9 Hz, 1H), 4.62 (t, *J* = 3.7 Hz, 1H), 4.23 (dd, *J* = 11.9, 6.4 Hz, 1H), 4.07 (dd, *J* = 6.1, 3.0 Hz, 2H), 4.02 (dd, *J* = 12.0, 7.4 Hz, 1H), 3.93 (s, 2H), 3.92 – 3.86 (m, 1H), 3.54 – 3.47 (m, 1H), 2.40 (t, *J* = 2.5 Hz, 1H), 2.17 – 2.08 (m, 4H), 2.07 – 1.98 (m, 4H), 1.89 – 1.79 (m, 1H), 1.75 – 1.70 (m, 1H), 1.68 (s, 3H), 1.65 (s, 3H), 1.60 (s, 3H), 1.58 – 1.46 (m, 4H).

### (2E,6E,10E)-3,7,11-trimethyl-12-(prop-2-yn-1-yloxy)dodeca-2,6,10-trien-1-ol (6)

To a solution of THP-protected alcohol **5** (1.00 g, 2.81 mmol, 1.0 equiv) in EtOH (8 mL), PPTs (212 mg, 0.85 mmol, 0.3 equiv) was added at RT. The reaction mixture was then heated to 60 °C for 3 h. After the consumption of starting materials (monitored by TLC), the solvent was evaporated and purified by column chromatography to afford 640 mg (82% yield) of the desired product as a pale yellow oil. The average reaction yield when performed 5 times was 81%. That data is summarized in Table S3. ^1^H NMR (500 MHz, CDCl3) δ 5.44 (t, *J* = 6.4 Hz, 2H), 5.14 (t, *J* = 6.6 Hz, 1H), 4.18 (d, *J* = 6.9 Hz, 2H), 4.10 (d, *J* = 2.3 Hz, 2H), 3.96 (s, 2H), 2.43 (t, *J* = 2.1 Hz, 1H), 2.14 (dt, *J* = 14.3, 6.8 Hz, 4H), 2.06 (dd, *J* = 14.3, 6.6 Hz, 4H), 1.71 (s, 3H), 1.67 (s, 3H), 1.63 (s, 3H). (SM-1028, SM-1013).

### (2E,6E,10E)-3,7,11-trimethyl-12-(prop-2-yn-1-yloxy)dodeca-2,6,10-trien-1-yl diphosphate (1)

PPh3 (polymer-supported beads, 208 mg, 0.80 mmol, 2.5 equiv) was suspended in 4 mL CH2Cl2 and the reaction mixture was stirred until all the beads were swelled (minimum 45 min) at RT under a N2 atmosphere. Next, a solution of alcohol **6** (100 mg, 0.32 mmol, 1.0 equiv) in CH2Cl2 was added to the suspension and stirred at RT. After 15 min, CBr4 (170 mg, 0.48 mmol, 1.5 equiv) was added to the reaction mixture that was then stirred for 3 h under N2 at RT. After completion (monitored by TLC), the beads were removed by filtration (using filter paper) to yield crude bromide **7** (crude product, 130 mg) that was then used without purification. Hence, crude **7** was dissolved in dry CH3CN (2.0 mL) and added into the reaction mixture containing (*n*-Bu4N)3HP2O7 (560 mg, 0.64 mmol, 2.0 equiv) in a glass vial and stirred at RT. After 4 h of reaction, the reaction mixture was passed through Dowex resin (Amberchrome® 50WX8 100-200H, hydrogen form, 50-100 mesh) ion-exchange to convert the product to its ammonium salt. For that process, resin was packed into a column, washed with 3 bed volumes of H2O/conc. NH4OH (3/1, v/v), and equilibrated with 4 bed volumes of 25 mM NH4HCO3/*i*-PrOH (98/2, v/v, solvent C). The crude product was loaded onto the ion-exchange column and eluted with 100 mL of solvent C; 20 mL fractions were collected. Fractions containing the desired product (determined by ESI MS) were pooled and lyophilized. Next the crude material was suspended in 3 mL 0.1 M NH4HCO3 and 12 mL 1:1 CH3CN:*i*-PrOH was added to the solution which was then vortexed and then centrifuged for 3 min. The resulting supernatant was decanted and the precipitate was extracted a second time with (1:1 CH3CN: *i*-PrOH) followed by centrifugation. The combined supernatant fractions were evaporated, and the resulting crude material was then purified via cellulose column chromatography (Cellulose fibers, (medium), Sigma-Aldrich C6288). A column (2.5 x 18 cm) was packed and eluted with 90:10 THF: 0.1 M NH4HCO3. 200 mL of 90:10 THF:0.1M NH4HCO3 was used first followed by 200 mL 80:20 THF:0.1 M NH4HCO3 and then 100 mL 70:30 THF:0.1 M NH4HCO3. This was carried out at a flow rate of 5.0-6.0 mL/min). Fractions (8 mL) were collected in 13 x 100 mm test tubes and checked by ESI-MS for the presence of the product ([M-H]^-^, m/z = 435.1, negative ion mode). The pure fractions containing the diphosphate were pooled and the THF was first removed by rotary evaporation followed by lyophilization to yield 39 mg (25% yield) of a white powder. In this case, the yield of diphosphate product was determined by the ^31^P NMR spectroscopy using an internal standard (Na2HPO4) in D2O. This allowed solutions of known concentration of the diphosphate to be prepared for subsequent biological experiments. The average reaction yield when performed 7 times was 25%. That data is summarized in Table S4. ^1^H NMR (500 MHz, D2O) δ 5.34 – 5.16 (m, 1H), 5.00 (t, *J* = 6.8 Hz, 1H), 4.30 (t, *J* = 6.3 Hz, 2H), 3.87 (s, 2H), 3.75 (s, 2H), 2.01 – 1.79 (m, 8H), 1.56 (s, 3H), 1.45 (s, 6H); ^13^C NMR (101 MHz, D2O) δ 141.9, 134.7, 130.9, 129.5, 124.7, 120.2, 75.3, 62.3, 55.6, 39.34, 38.9, 30.2, 26.3, 26.0, 25.4, 15.9, 15.6, 13.6; ^31^P NMR (162 MHz, D2O) δ -6.41 (d, *J* = 23.0 Hz, 1P), -10.46 (d, *J* = 22.8 Hz, 1P).

### Maintenance of mouse ESCs

The mouse ES cell line expressing GFP and CD2 under the control of the Hb9 promoter was described elsewhere.^21,22^ Mouse ES cell lines were maintained on gelatinized tissue culture plastic on a monolayer of irradiated mouse embryonic fibroblasts (MEFs) (GlobalStem) originally seeded at a density of 15,000 MEFs per cm^2^, in DMEM medium (Millipore) supplemented with penicillin/streptomycin (P/S; 100 Units/ml and 100 μg/ml, respectively; Millipore), 2 mM L-glutamine (Gibco), 100 μM non-essential amino acids (Gibco), 1% nucleosides (Millipore), 10% FBS (Millipore) and LIF (Millipore, ESG1106). Cells were maintained at 37°C, 5% CO2, and regularly passaged using 0.5% trypsin (Millipore).

### Mouse ESC differentiation into motor neurons

Mouse ESCs were differentiated into MNs using a well-established differentiation protocol.^21,22^ Briefly, the ESC line Hb9::eGFP-CD2 was plated on gelatinized 25 cm^2^ flasks in DMEM media (Millipore Sigma) containing 15% ES-FBS (Millipore Sigma), 1% penicillin/streptomycin (Gibco), 1% glutamine (Gibco), 1% non-essential amino acids (Millipore Sigma), 1% nucleosides (Millipore Sigma), 1% β-mercaptoethanol (Millipore Sigma), 1% sodium pyruvate (Sigma-Aldrich) and 0.1% LIF (Millipore Sigma). After 48 h, the cells were trypsinized and plated at 1 million cells in a 10 cm^2^ culture dish (BD Falcon) containing αDFNK media containing a 1:1 ratio of Advanced DMEM (Gibco) and Neurobasal A (Thermofisher Scientific), 10% Knock Out Serum Replacement (Millipore Sigma), 1% penicillin/streptomycin (Gibco), 1% L-glutamine (Gibco) and 1% β-mercaptoethanol (Millipore). After 24 h, the supernatant containing embryoid bodies (EBs) was removed, gently pelleted at 160 g for 1 min, and transferred to an ultra-low attachment dish (Corning). The next day, differentiating factors 1 µM retinoic acid (Sigma-Aldrich) and 0.25 µM Smoothened Agonist (Calbiochem) were added for 72 h. Differentiated EBs were monitored for differentiation based on GFP expression under live fluorescence microscopy.

### Mouse ESC differentiation into astrocytes

Mouse ESCs were differentiated into astrocytes using a combination and adjustment of two different protocols.^39,40^ Briefly, 1 million ESCs were seeded onto ultra-low attachment dishes (Corning) and suspended in 10 mL of ADFNK media (Advanced DMEM/F-12:Neurobasal medium (1:1) supplemented with 10% Knockout Serum Replacement, 1% L-Glutamine, 100 μM β-mercaptoethanol, and 1% Pen/Strep). After 2 days, single cells formed embryoid bodies (EBs). EBs were exposed to fresh ADFNK media containing 1 μM retinoic acid (RA) (Sigma Aldrich) on day 2. On day 5, RA was removed, and fresh ADFNK media was added. On day 7, EBs were dissociated using 0.5% trypsin EDTA (Gibco) for 5 min at 37 °C. After halting the trypsinization with horse serum (Thermofisher Scientific), the EBs were gently triturated using a 1 mL tip and transferred to a 4% BSA cushion, spun at 300 × *g* for 5 min. Single cells were seeded onto 100 μg/mL poly-L-ornithine (Sigma-Aldrich)-coated surfaces in DMEM (Gibco) supplemented with 10% FBS, 2 mM L-glutamine (Gibco), 100 Units/mL penicillin (Gibco), 100 μg/mL streptomycin (Gibco). Cells were passed weekly. The cells were assessed via bright field microscopy every day and cultured for up to 28 days. The presence of astrocytes was detected by immunofluorescence.

### Metabolic labeling of ES-MNs and ES-As and their ESC progenitors

Cultured cells were labeled following a modified version of a well-established protocol for metabolic labeling of prenylated proteins.^41^ Briefly, ES-MNs were differentiated from the Hb9::eGFP-CD2 ESC line as described above. On day 3 of differentiation, EBs were pre-treated with 10 µM lovastatin (Cayman Chemical) for 6 h or 2 µM lovastatin for 24 h, followed by incubation with FPP (25 µM) or C15AlkOPP (25 µM) for 48 h or 72 h. On day 3 of differentiation, labelled EB-MNs were dissociated using 0.5% trypsin EDTA (Gibco) for 5 min at 37 °C. After halting the trypsinization with horse serum (Thermofisher Scientific), the EB-MNs were gently triturated using a 1 mL tip and transferred to a 4% BSA cushion, spun at 300 × *g* for 5 min. The pellet was resuspended in L15 medium (Sigma) and purified by magnetic cell sorting as described in Ikiz *et al.*^22^ Briefly, cell suspensions were incubated for 20 min at 4°C in 80 µL of L15 medium containing 2.1 µg of anti-rat CD2 antibody (Invitrogen) per 15 million dissociated cells. After washing with L15 medium, cells were incubated for 20 min at 4°C with an anti-mouse antibody conjugated to magnetic microbeads (Miltenyi Biotec). Finally, cells were passed through a magnetic column to separate CD2-GFP^+^ cells from the other types. Post-sorting, the cells were collected for proteomic analysis of prenylated proteins.

After differentiation from ESCs, ES-As were grown in 100-mm dishes containing 10 mL of DMEM (Gibco) media with 10% FBS (Gibco), and 1% penicillin-streptomycin (Gibco) two days prior to treatment. The media was removed and replaced with 5 mL of fresh media, followed by pre-treatment with 10 μM lovastatin for 24 h. After pre-treatment, the media was replaced to remove the lovastatin, followed by the addition of 25 μM C15AlkOPP or FPP. After 72 h incubation, the cells were collected for proteomic analysis of prenylated proteins.

ESCs were grown in gelatin-coated 100-mm dishes containing 10 mL of ES medium (see above) two days prior to treatment. The media was removed and replaced with 5 mL of fresh media, followed by pre-treatment with 10 μM lovastatin for 24 h. After pre-treatment, the media was replaced to remove the lovastatin, followed by the addition of 25 μM C15AlkOPP or FPP. After 72 h incubation, the cells were collected for proteomic analysis of prenylated proteins.

### Gel-based analysis of metabolic labeling samples

Labeled proteins were analyzed by in-gel fluorescence as described previously.^15^ Briefly, labeled cell samples were lysed in 1X PBS containing 1% SDS, 2.4 *μ*M PMSF, 200 units/nL benzonase nuclease (Sigma-Aldrich), protease inhibitor cocktail *via* sonication. BCA assay (Thermo Fisher Scientific) was then used to quantify protein yields and aliquots of 100 *μ*g were subjected to click reaction with TAMRA-N3 (25 *μ*M TAMRA-N3 (BroadPharm), 1 mM TCEP, 0.1 mM TBTA (Sigma-Aldrich), and 1 mM CuSO4) for 1 hour under ambient conditions. Proteins were precipitated out using ProteoExtract precipitation kit (Calbiochem) and dissolved in 1X Laemmli buffer. Protein samples were resolved in 12% SDS-PAGE gels to detect TAMRA fluorescence using Typhoon FLA 9500 (GE Healthcare) and images were processed and formatted using ImageJ.

### Chemical proteomic analysis

Enrichment and subsequent LC-MS analysis were performed as previously described.^15^ Briefly, protein lysates (2 mg in 1 mL) from cells treated with lovastatin and C15AlkOPP or FPP under optimized conditions were subjected to click reaction with biotin-N3 (100 *μ* M biotin-N3 (BroadPharm), 1 mM TCEP, 0.1 mM TBTA, and 1 mM CuSO4) for 1.5 hours under ambient conditions. Proteins were precipitated out using chloroform-methanol-water (1:4:3) pelleted by centrifugation. Pellets were dissolved in 1 mL of 1X PBS + 1% SDS and quantified using a BCA assay; concentrations were normalized across all samples. Samples were prepared as 1 mg in 500 *μ*L and incubated with 100 *μ*L (settled resin) of pre-washed Neutravidin® agarose beads (Thermo Scientific) for 1.5 hrs. Washings were performed as follows: 1X PBS + 1% SDS (3x), 1X PBS (1x), 8 M urea in 50 mM TEAB (3x), and 50 mM TEAB (3x) with 1 mL per wash. Tryptic digestion was performed on-bead with 1.5 μg of sequencing grade trypsin (Promega Corp.) overnight at 37°C. Peptides were collected in 50 mM TEAB, dried dissolved in 100 mM TEAB and quantified using BCA assay. TMT-labeling was performed in 10 μg of samples supplemented with 150 fmol of yADH1 following the manufacturer’s protocol. Samples were fractionated on SDB-XC extraction disks (3M, 1.07 mm x 0.50 mm i.d.) under high pH reversed phase conditions into 7 fractions (5,10, 15, 20, 22.5, 27.5, and 80% acetonitrile in 200 mM ammonium formate pH 10). Fractions 5 and 10% were combined and each sample were dried and dissolved in 0.1% formic acid in water.

Peptide samples were analyzed in a RSLC Ultimate 3000 nano-UHPLC (Dionex) with a reversed-phase column (75 *μ*m i.d., 45 cm) containing ProntoSIL C18AQ 3 *μ*m media at a flow rate of 300 nL/min and sprayed into the Orbitrap using a Nanospray Flex source (Thermo Fisher Scientific). An MS2-based quantitative proteomic analysis was performed using an Orbitrap Fusion instrument (Thermo Fisher Scientific) equipped with RSLC Ultimate 3000 nano-UHPLC. A data-dependent acquisition was performed for MS1 scans under 60,000 resolution in a 320-2000 *m/z* range, with a normalized AGC target at 125% and a max IT of 50 ms. Fragmentation of precursors was performed using HCD at NCE of 38% with a 1.5 *m/z* isolation window, a standard AGC target, and a max IT of 200 ms at 15,000 resolution.

Database searches were performed using Andromeda embedded in MaxQuant (version 1.6.2.10) against the non-redundant human (UP000005640) or mouse (UP0000000589) databases from Uniprot (EMBL-EBI, April 2018 release). A full tryptic digestion was assumed with allowed missed cleavages up to 3. Fixed modifications were set for the TMT labels on both the N-terminal and lysine and variable modifications for methionine oxidation and N-term acetylation. Statistical analyses were performed in Perseus (version 1.6.0.7) Proteins that were identified only by site, potential contaminant, or reversed were removed. Only proteins with 3 out of 6 valid values across the six TMT channels were pursued. Missing values were imputed based on a normal distribution, and reporter ion values were normalized by subtracting rows by means and columns by median. A two-sample t-test with FDR = 0.01 and s0 = 0.5 was used for statistical analysis and generation of volcano plots. Further data processing was performed using Excel.

### Transcriptomic versus enrichment analysis

The list of all prenylated proteins found in the ES-MNs was ungrouped; for each protein in the group, the same Log 2 (fold change) and -Log 10 (P-value) were used. This list was cross-referenced with a previously collected transcriptomic analysis of the ES-derived motor neurons.^22^ From the list of proteins, all but two proteins were found to be significant, and one protein was not found in the transcriptomics dataset; these were not included in the comparative analysis. The transcript counts of the common proteins were normalized by gene lengths determined using the Ensembl gene bank. From there, the counts were then normalized by the smallest value, Eras. Then, the normalized counts were transformed using a Log 2 scale. The correlation of these data sets was determined using a Spearman’s rank-order correlation with a one-tail distribution.

### Bioinformatic analysis

The log fold change values, scaled per row, of the prenylated proteins shared across the three cell type populations ES, Es-As, and ES-MN were visualized in a heatmap performed using the heatmap.2 function of the gplots R package. In the heatmap, the hierarchical clustering was performed between rows. The ORA (Over Representation Analysis) of the three protein signatures was performed using the R package gprofiler2 (version 0.2.1), The PPI (Protein-Protein Interactions) was performed with STRING (Version 12.0).

## Data Availability

The raw mass spectrometry files for the proteomics data have been deposited to the ProteomeXchange Consortium via the PRIDE partner repository with the dataset identifier PXD040354.

## Supporting Information

Supporting information including summary of yields from duplicate reactions, analysis of oxidation reaction including ESI-MS and NMR data, and spectra for all synthetic intermediates and final product **1**. Complete lists of prenylated proteins and gprofiler analysis are also included. This material is available free of charge via the internet.

## Acknowledgements

The authors acknowledge the Minnesota Supercomputing Institute (MSI) at the University of Minnesota for providing resources that contributed to the prenylomic research results reported within this paper (http://www.msi.umn.edu) and Y. Zhao and P. Villalta for the assistance with proteomic data collection in the Analytical Biochemistry Shared Resource of the Masonic Cancer Center, designated by the National Cancer Institute and supported by P30 CA077598.

## References

(1) Zhang, F. L.; Casey, P. J. Protein Prenylation: Molecular Mechanisms and Functional Consequences. Annu. Rev. Biochem. 1996, 65 (1), 241–269.

(2) Marshall, C. J. Protein Prenylation: A Mediator of Protein-Protein Interactions. Science 1993, 259 (5103), 1865–1866. 10.1126/science.8456312.

(3) Palsuledesai, C. C.; Distefano, M. D. Protein Prenylation: Enzymes, Therapeutics, and Biotechnology Applications. ACS Chem. Biol. 2015, 10 (1), 51–62. 10.1021/cb500791f.

4. Slater, H. FDA Grants Fast Track Designation to Tipifarnib for the Treatment of Patients with HNSCC https://www.cancernetwork.com/view/fda-grants-fast-track-designation-tipifarnib-treatment-patients-hnscc (accessed Feb 3, 2024).

(5) Wang, Y.; Xu, F.; Nichols, C. B.; Shi, Y.; Hellinga, H. W.; Alspaugh, J. A.; Distefano, M. D.; Beese, L. S. Structure-Guided Discovery of Potent Antifungals That Prevent Ras Signaling by Inhibiting Protein Farnesyltransferase. J. Med. Chem. 2022, 65 (20), 13753–13770. 10.1021/acs.jmedchem.2c00902.

6. FDA Approves First Treatment for Hutchinson-Gilford Progeria Syndrome and Some Progeroid Laminopathies https://www.fda.gov/news-events/press-announcements/fda-approves-first-treatment-hutchinson-gilford-progeria-syndrome-and-some-progeroid-laminopathies.

(7) Morgan, M. A.; Ganser, A.; Reuter, C. W. M. Therapeutic Efficacy of Prenylation Inhibitors in the Treatment of Myeloid Leukemia. Leukemia 2003, 17 (8), 1482–1498. 10.1038/sj.leu.2403024.

(8) Jeong, A.; Suazo, K. F.; Wood, W. G.; Distefano, M. D.; Li, L. Isoprenoids and Protein Prenylation: Implications in the Pathogenesis and Therapeutic Intervention of Alzheimer’s Disease. Crit. Rev. Biochem. Mol. Biol. 2018, 53 (3), 279–310.

(9) Li, H.; Kuwajima, T.; Oakley, D.; Nikulina, E.; Hou, J.; Yang, W. S.; Lowry, E. R.; Lamas, N. J.; Amoroso, M. W.; Croft, G. F.;, et al. Protein Prenylation Constitutes an Endogenous Brake on Axonal Growth. Cell Rep. 2016, 16 (2), 545–558.

(10) Charron, G.; Li, M. M. H.; MacDonald, M. R.; Hang, H. C. Prenylome Profiling Reveals S-Farnesylation Is Crucial for Membrane Targeting and Antiviral Activity of ZAP Long-Isoform. Proc. Natl. Acad. Sci. 2013, 110 (27), 11085–11090.

(11) Storck, E. M.; Morales-Sanfrutos, J.; Serwa, R. A.; Panyain, N.; Lanyon-Hogg, T.; Tolmachova, T.; Ventimiglia, L. N.; Martin-Serrano, J.; Seabra, M. C.; Wojciak-Stothard, B.;, et al. Dual Chemical Probes Enable Quantitative System-Wide Analysis of Protein Prenylation and Prenylation Dynamics. Nat. Chem. 2019, 11 (6), 552–561. 10.1038/s41557-019-0237-6.

(12) Suazo, K. F.; Schaber, C.; Palsuledesai, C. C.; Odom John, A. R.; Distefano, M. D.; Feachem, R. G. A.; Ashley, E. A.; Seidlein, L. Von; Dondorp, A.; Eastman, R. T.;, et al. Global Proteomic Analysis of Prenylated Proteins in Plasmodium Falciparum Using an Alkyne-Modified Isoprenoid Analogue. Sci. Rep. 2016, 6, 38615. 10.1038/srep38615.

(13) Palsuledesai, C. C.; Ochocki, J. D.; Kuhns, M. M.; Wang, Y.-C.; Warmka, J. K.; Chernick, D. S.; Wattenberg, E. V; Li, L.; Arriaga, E. A.; Distefano, M. D. Metabolic Labeling with an Alkyne-Modified Isoprenoid Analog Facilitates Imaging and Quantification of the Prenylome in Cells. ACS Chem. Biol. 2016, 11 (10), 2820–2828. 10.1021/acschembio.6b00421.

(14) Maxwell, Z. A.; Suazo, K. F.; Brown, H. M. G.; Distefano, M. D.; Arriaga, E. A. Combining Isoprenoid Probes with Antibody Markers for Mass Cytometric Analysis of Prenylation in Single Cells. Anal. Chem. 2022, 94 (33), 11521–11528. 10.1021/acs.analchem.2c01509.

(15) Suazo, K. F.; Jeong, A.; Ahmadi, M.; Brown, C.; Qu, W.; Li, L.; Distefano, M. D. Metabolic Labeling with an Alkyne Probe Reveals Similarities and Differences in the Prenylomes of Several Brain-Derived Cell Lines and Primary Cells. Sci. Rep. 2021, 11 (1), 4367. 10.1038/s41598-021-83666-3.

(16) Jeong, A.; Auger, S. A.; Maity, S.; Li, L.; Distefano, M. D. In Vivo Prenylomic Profiling in the Brain of a Transgenic Mouse Model of Alzheimer’s Disease Reveals Increased Prenylation of a Key Set of Proteins. bioRxiv 2022, 2022.04.01.486487. 10.1101/2022.04.01.486487.

(17) Petenkova, A.; Auger, S. A.; Lamb, J.; Quellier, D.; Carter, C.; To, O. T.; Milosevic, J.; Barghout, R.; Kugadas, A.; Lu, X.;, et al. Prenylcysteine Oxidase 1 like Protein Is Required for Neutrophil Bactericidal Activities. Nat. Commun. 2023, 14 (1), 2761. 10.1038/s41467-023-38447-z.

(18) Hosokawa, A.; Wollack, J. W.; Zhang, Z.; Chen, L.; Barany, G.; Distefano, M. D. Evaluation of an Alkyne-Containing Analogue of Farnesyl Diphosphate as a Dual Substrate for Protein-Prenyltransferases. Int. J. Pept. Res. Ther. 2007, 13 (1), 345–354.

(19) Nagai, M.; Re, D. B.; Nagata, T.; Chalazonitis, A.; Jessell, T. M.; Wichterle, H.; Przedborski, S. Astrocytes Expressing ALS-Linked Mutated SOD1 Release Factors Selectively Toxic to Motor Neurons. Nat. Neurosci. 2007, 10 (5), 615–622. 10.1038/nn1876.

(20) Wichterle, H.; Peljto, M. Differentiation of Mouse Embryonic Stem Cells to Spinal Motor Neurons. Curr. Protoc. Stem Cell Biol. 2008, 5 (1), 1H.1.1–1H.1.9. 10.1002/9780470151808.sc01h01s5.

(21) Wichterle, H.; Lieberam, I.; Porter, J. A.; Jessell, T. M. Directed Differentiation of Embryonic Stem Cells into Motor Neurons. Cell 2002, 110 (3), 385–397. 10.1016/S0092-8674(02)00835-8.

(22) Ikiz, B.; Alvarez, M. J.; Ré, D. B.; Le Verche, V.; Politi, K.; Lotti, F.; Phani, S.; Pradhan, R.; Yu, C.; Croft, G. F.;, et al. The Regulatory Machinery of Neurodegeneration in In Vitro Models of Amyotrophic Lateral Sclerosis. Cell Rep. 2015, 12 (2), 335–345. 10.1016/j.celrep.2015.06.019.

(23) Goldstein, J. L.; Brown, M. S. Regulation of the Mevalonate Pathway. Nature 1990, 343, 425.

(24) Ahmadi, M.; Suazo, K. F.; Distefano, M. D. Optimization of Metabolic Labeling with Alkyne-Containing Isoprenoid Probes BT - Protein Lipidation: Methods and Protocols; Linder, M. E., Ed.; Springer New York: New York, NY, 2019; pp 35–43.

(25) Maurer-Stroh, S.; Eisenhaber, F. Refinement and Prediction of Protein Prenylation Motifs. Genome Biol. 2005, 6 (6), 1–15. 10.1186/gb-2005-6-6-r55.

(26) Kiral, F. R.; Kohrs, F. E.; Jin, E. J.; Hiesinger, P. R. Rab GTPases and Membrane Trafficking In Neurodegeneration. Curr. Biol. 2018, 28 (8), R471–R486. 10.1016/j.cub.2018.02.010.

(27) Kho, Y.; Kim, S. C.; Jiang, C.; Barma, D.; Kwon, S. W.; Cheng, J.; Jaunbergs, J.; Weinbaum, C.; Tamanoi, F.; Falck, J.;, et al. A Tagging-via-Substrate Technology for Detection and Proteomics of Farnesylated Proteins. Proc. Natl. Acad. Sci. U. S. A. 2004, 101 (34), 12479–12484. 10.1073/pnas.0403413101.

(28) Palsuledesai, C. C.; Ochocki, J. D.; Markowski, T. W.; Distefano, M. D. A Combination of Metabolic Labeling and 2D-DIGE Analysis in Response to a Farnesyltransferase Inhibitor Facilitates the Discovery of New Prenylated Proteins. Mol. BioSyst. 2014, 10 (5), 1094–1103. 10.1039/C3MB70593E.

(29) Suazo, K. F.; Park, K.-Y.; Distefano, M. D. A Not-So-Ancient Grease History: Click Chemistry and Protein Lipid Modifications. Chem. Rev. 2021, 121 (12), 7178–7248. 10.1021/acs.chemrev.0c01108.

(30) Osmani, N.; Peglion, F.; Chavrier, P.; Etienne-Manneville, S. Cdc42 Localization and Cell Polarity Depend on Membrane Traffic. J. Cell Biol. 2010, 191 (7), 1261–1269. 10.1083/jcb.201003091.

(31) Wang, Y.; Tang, B. L. SNAREs in Neurons – beyond Synaptic Vesicle Exocytosis (Review). Mol. Membr. Biol. 2006, 23 (5), 377–384. 10.1080/09687860600776734.

(32) Dai, Y.; Seeger, M.; Weng, J.; Song, S.; Wang, W.; Tan, Y.-W. Lipid Regulated Intramolecular Conformational Dynamics of SNARE-Protein Ykt6. Sci. Rep. 2016, 6 (1), 30282. 10.1038/srep30282.

(33) Kawauchi, T.; Sekine, K.; Shikanai, M.; Chihama, K.; Tomita, K.; Kubo, K.; Nakajima, K.; Nabeshima, Y.; Hoshino, M. Rab GTPases-Dependent Endocytic Pathways Regulate Neuronal Migration and Maturation through N-Cadherin Trafficking. Neuron 2010, 67 (4), 588–602. 10.1016/j.neuron.2010.07.007.

(34) Nag, S.; Rani, S.; Mahanty, S.; Bissig, C.; Arora, P.; Azevedo, C.; Saiardi, A.; van der Sluijs, P.; Delevoye, C.; van Niel, G.;, et al. Rab4A Organizes Endosomal Domains for Sorting Cargo to Lysosome-Related Organelles. J. Cell Sci. 2018, 131 (18), jcs216226. 10.1242/jcs.216226.

(35) Lv, P.; Sheng, Y.; Zhao, Z.; Zhao, W.; Gu, L.; Xu, T.; Song, E. Targeted Disruption of Rab10 Causes Early Embryonic Lethality. Protein Cell 2015, 6 (6), 463–467. 10.1007/s13238-015-0150-8.

(36) Hart, C.; Chase, L. G.; Hajivandi, M.; Agnew, B. Metabolic Labeling and Click Chemistry Detection of Glycoprotein Markers of Mesenchymal Stem Cell Differentiation BT - Mesenchymal Stem Cell Assays and Applications; Vemuri, M., Chase, L. G., Rao, M. S., Eds.; Humana Press: Totowa, NJ, 2011; pp 459–484. 10.1007/978-1-60761-999-4_33.

(37) Konze, S. A.; Cajic, S.; Oberbeck, A.; Hennig, R.; Pich, A.; Rapp, E.; Buettner, F. F. R. Quantitative Assessment of Sialo-Glycoproteins and N-Glycans during Cardiomyogenic Differentiation of Human Induced Pluripotent Stem Cells. ChemBioChem 2017, 18 (13), 1317–1331. 10.1002/cbic.201700100.

(38) Bai, Q.-R.; Dong, L.; Hao, Y.; Chen, X.; Shen, Q. Metabolic Glycan Labeling-Assisted Discovery of Cell-Surface Markers for Primary Neural Stem and Progenitor Cells. Chem. Commun. 2018, 54 (43), 5486–5489. 10.1039/C8CC01535J.

(39) Roybon, L.; Lamas, N. J.; Garcia-Diaz, A.; Yang, E. J.; Sattler, R.; Jackson-Lewis, V.; Kim, Y. A.; Kachel, C. A.; Rothstein, J. D.; Przedborski, S.;, et al. Human Stem Cell-Derived Spinal Cord Astrocytes with Defined Mature or Reactive Phenotypes. Cell Rep. 2013, 4 (5), 1035–1048. 10.1016/j.celrep.2013.06.021.

(40) Kuegler, P. B.; Baumann, B. A.; Zimmer, B.; Keller, S.; Marx, A.; Kadereit, S.; Leist, M. GFAP-Independent Inflammatory Competence and Trophic Functions of Astrocytes Generated from Murine Embryonic Stem Cells. Glia 2012, 60 (2), 218–228. 10.1002/glia.21257.

(41) Suazo, K. F.; Hurben, A. K.; Liu, K.; Xu, F.; Thao, P.; Sudheer, C.; Li, L.; Distefano, M. Metabolic Labeling of Prenylated Proteins Using Alkyne-Modified Isoprenoid Analogues. Curr. Protoc. Chem. Biol. 2018, 10, e46. 10.1002/cpch.46.

