## Supporting Information for "Improved synthesis and application of an alkyne-functionalized isoprenoid analogue to study the prenylomes of motor neurons, astrocytes and their stem cell progenitors"

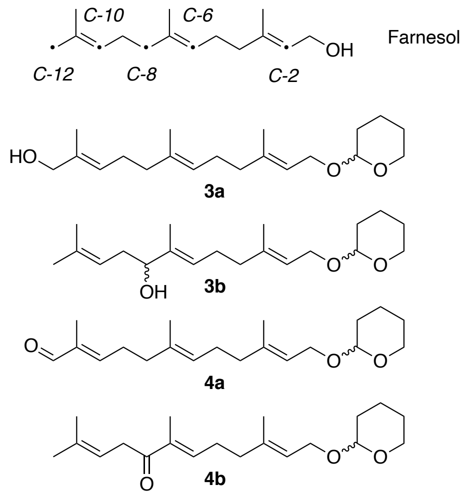

**Figure S1**. Products from the oxidation of compound **2**. The desired oxidation product **3a** containing a hydroxyl group at C-12 along with a second hydroxylated species at C-8 (**3b**). Carbonyl-containing products resulting from over-oxidation at C-12 (**4a**) and C-8 (**4b**) were also observed. The numbering scheme for Farnesol, used for all these products, is shown at the top.

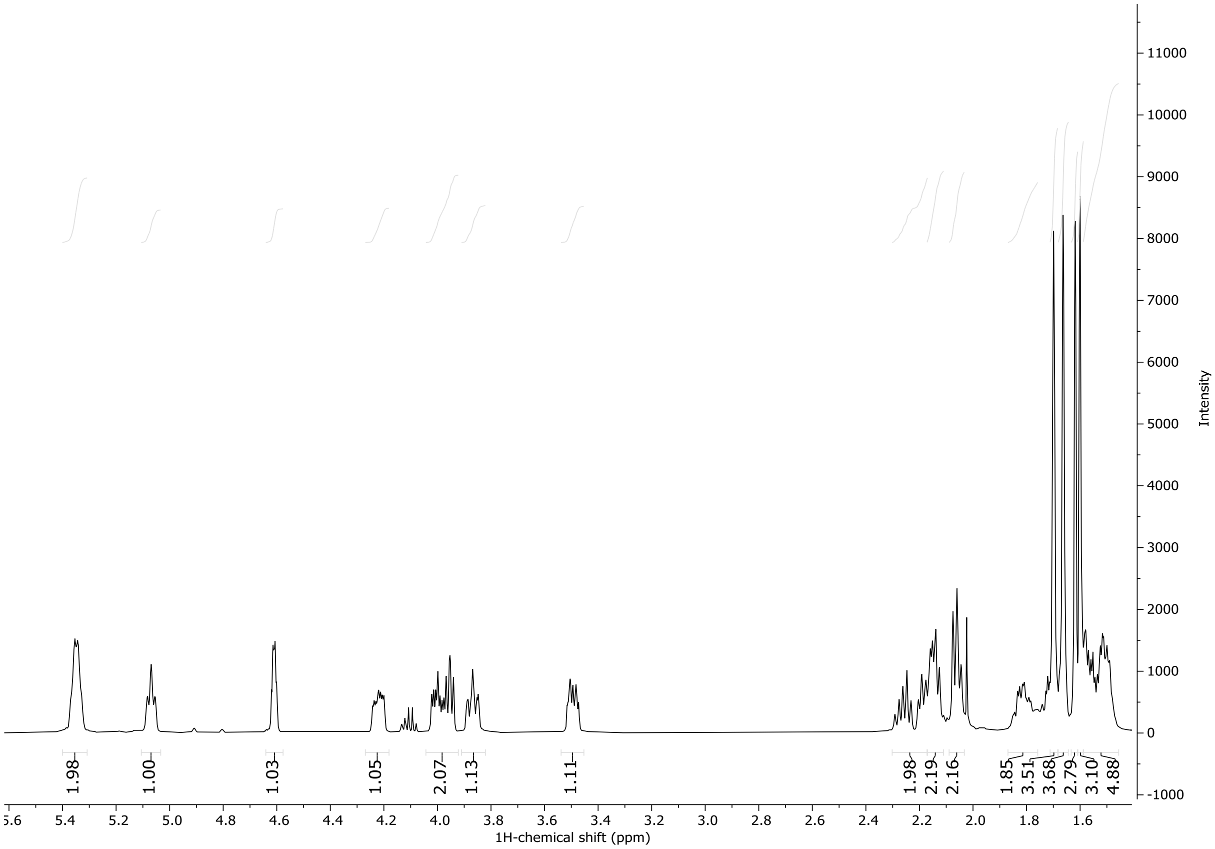

**Figure S2.** $^{1}$H-NMR analysis of **3b**.

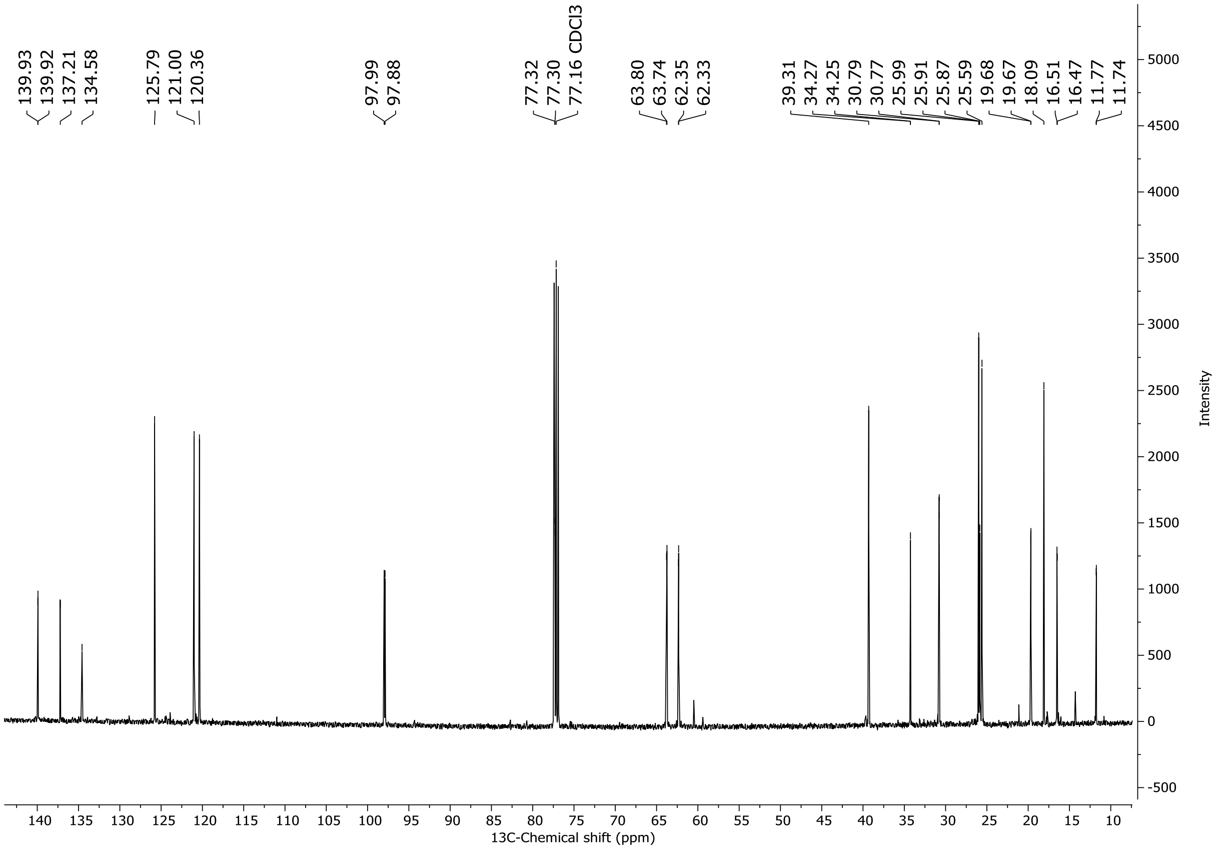

**Figure S3.** $^{13}$C-NMR analysis of **3b**.

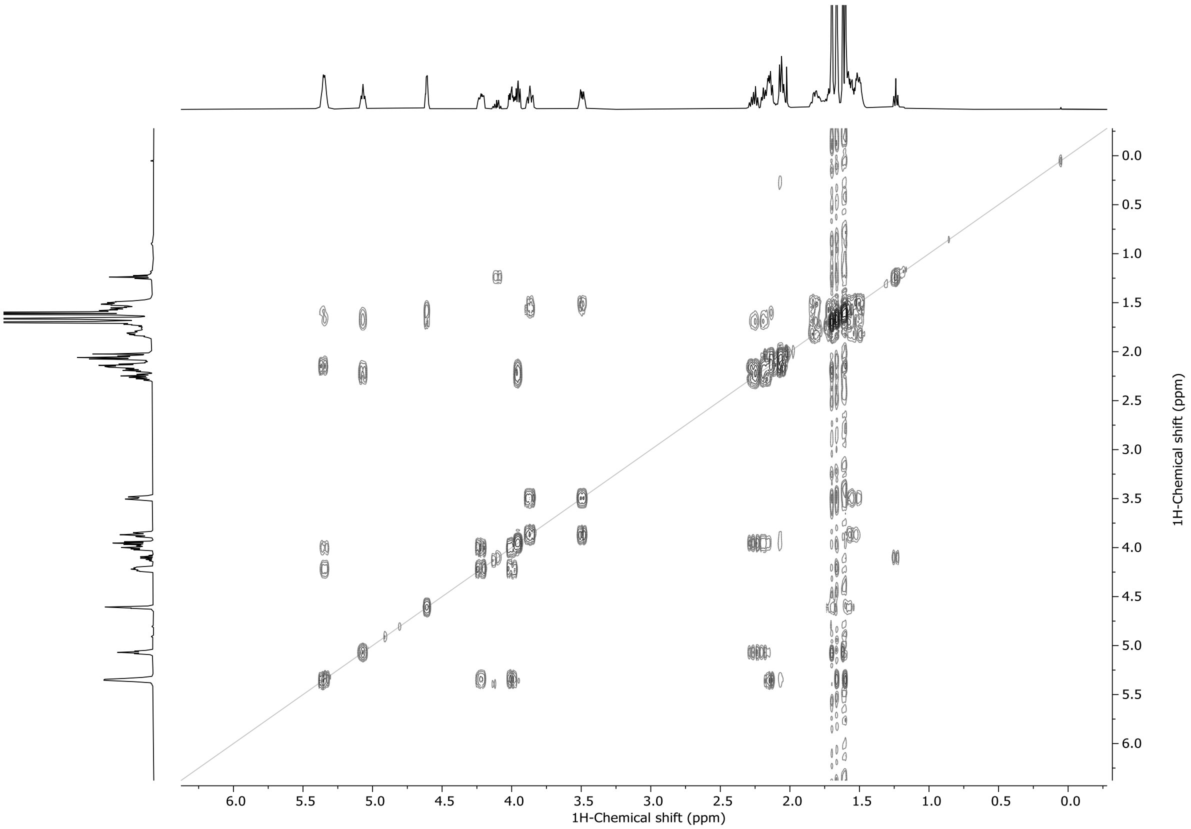

**Figure S4.** COSY NMR analysis of **3b**.

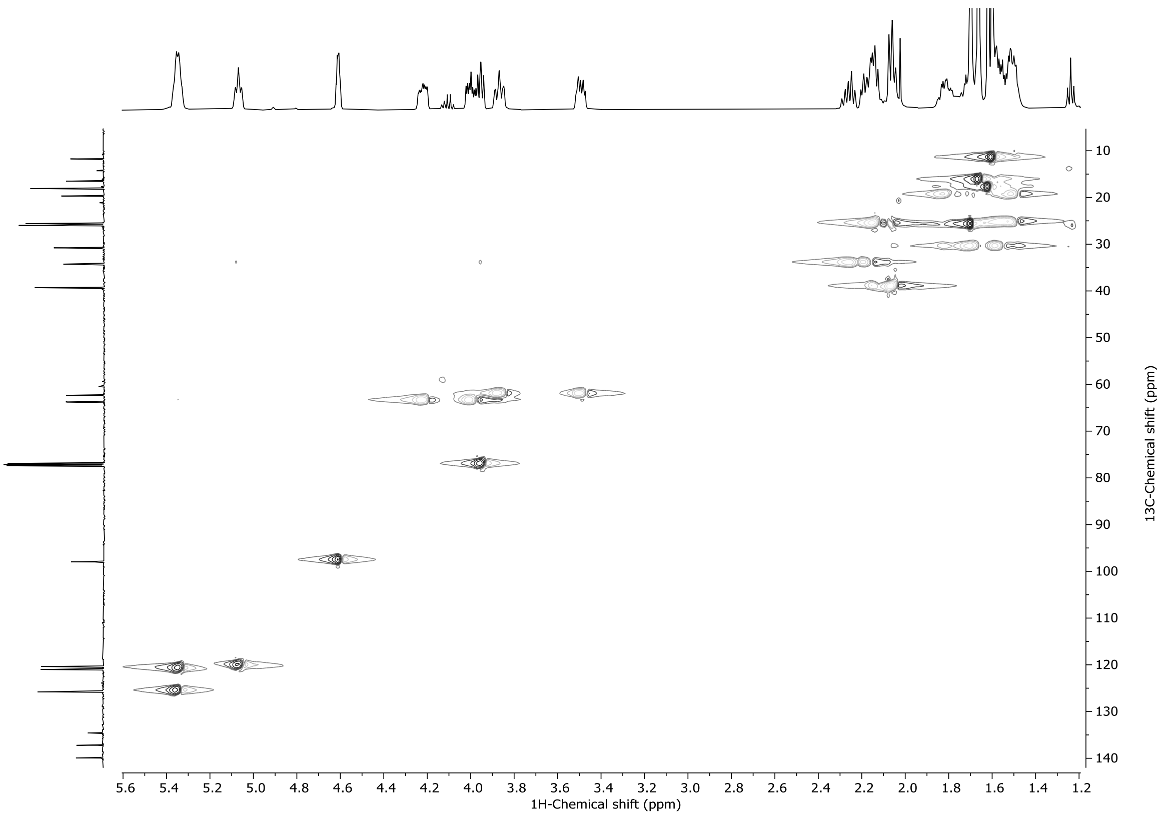

**Figure S5.** HSQC-NMR analysis of **3b**.

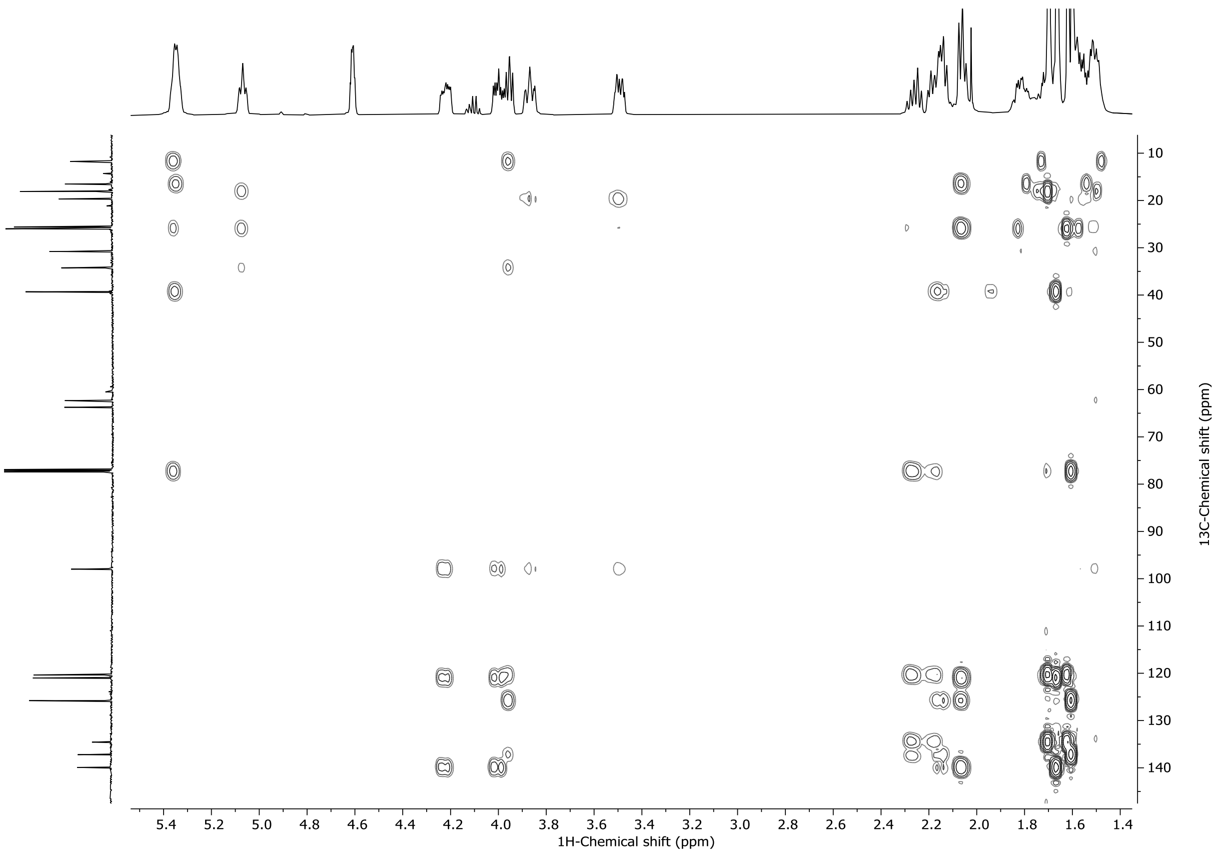

**Figure S6.** HMBC-NMR analysis of **3b**.

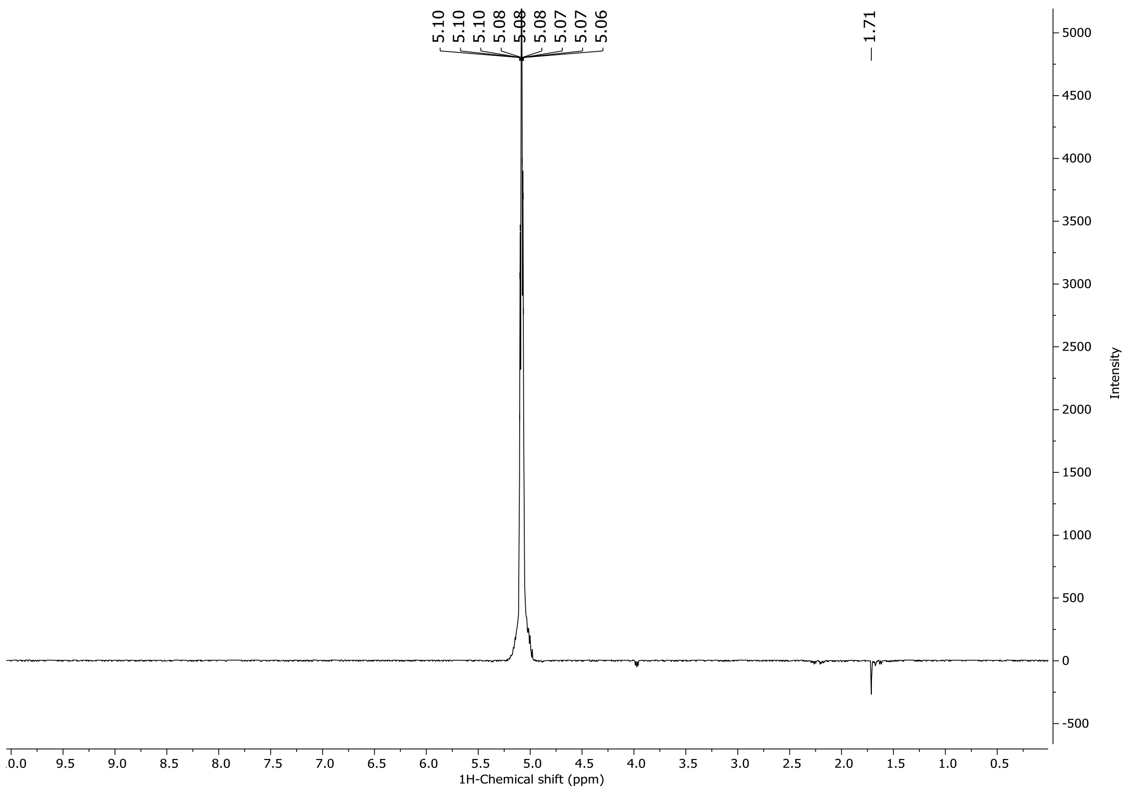

**Figure S7.** Selective Gradient NOESY analysis of **3b**.

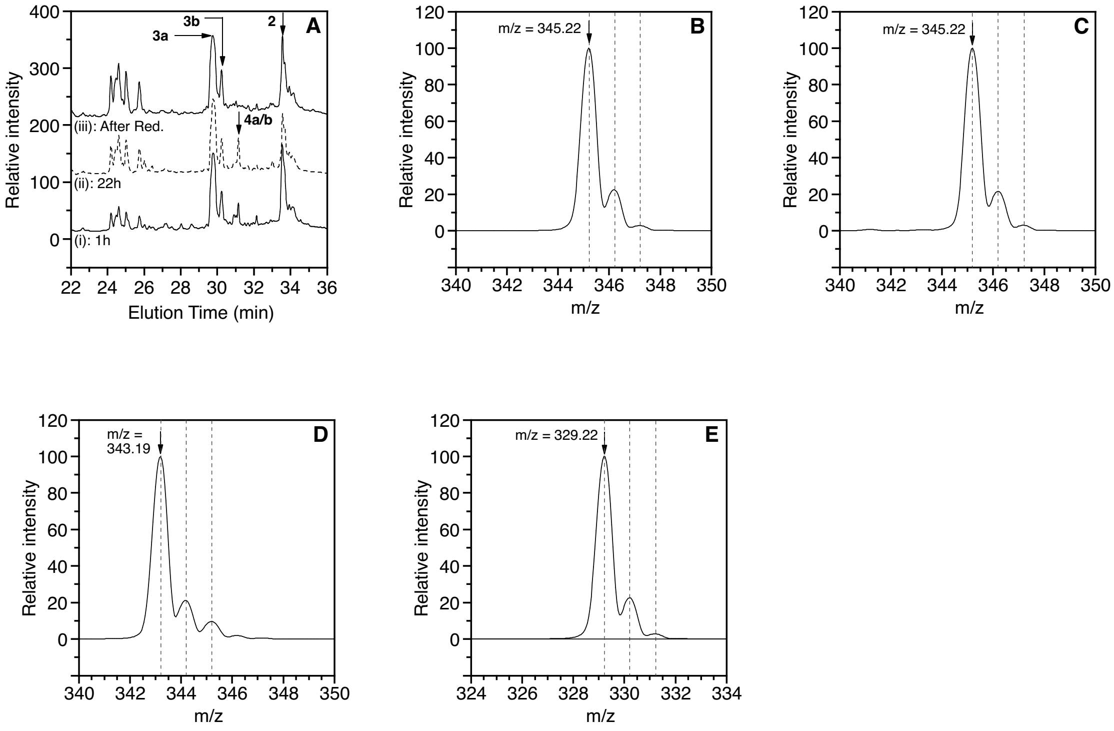

**Figure S8.** LC-MS analysis of the oxidation reaction of compound **2**. (A) LC-MS analysis of the oxidation reaction of compound **2** after 1 h (i), 22 h (ii) and after $NaBH_4$ reduction (iii). In the chromatograms, the peaks occurring at 29.7, 30.2, 31.2 and 33.6 are labeled as **3a**, **3b**, **4a** and/or **4b** and **2**, respectively. Those assignments were made on the basis of MS analysis and/or injection of the purified compounds. (B) MS analysis of the peak eluting at 29.7 min showing an m/z value of 345.22 compared to the calculated value of 345.24 for the $[M+Na]^+$ ion $C_{20}H_{34}O_3Na$. (C) MS analysis of the peak eluting at 30.2 min showing an m/z value of 345.22 compared to the calculated value of 345.24 for the $[M+Na]^+$ ion $C_{20}H_{34}O_3Na$. (D) MS analysis of the peak eluting at 31.2 min showing an m/z value of 343.19 compared to the calculated value of 343.23 for the $[M+Na]^+$ ion $C_{20}H_{32}O_3Na$. (E) MS analysis of the peak eluting at 33.6 min showing an m/z value of 329.22 compared to the calculated value of 229.25 for the $[M+Na]^+$ ion $C_{20}H_{34}O_2Na$. For each mass spectrum, the spacings between the parent peak and M+1 and M+2 isotopologues are highlighted with vertical dashed lines to illustrate that the observed species are singly charged in all cases.

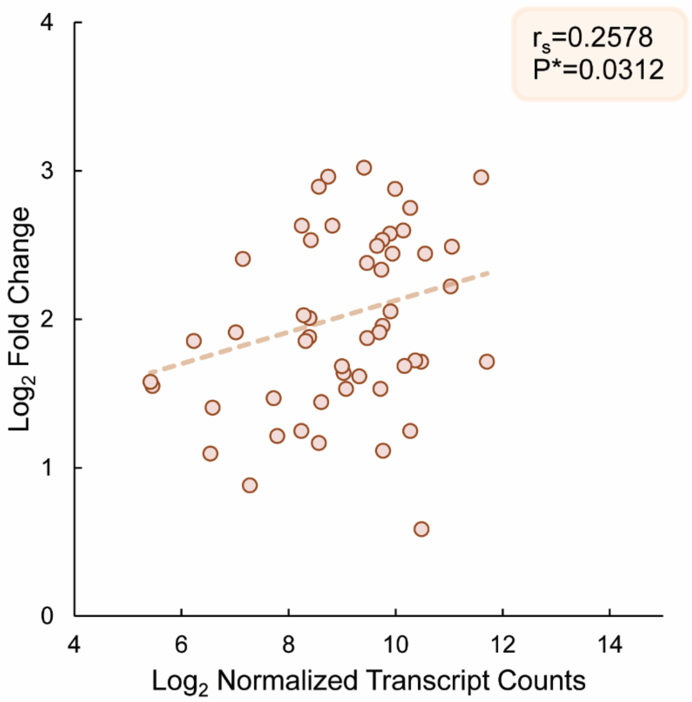

**Figure S9.** Correlation between normalized transcript counts and the fold-change of prenylated proteins ES-MNs. The $Log_2$ Fold change values of ungrouped prenylated proteins that were found in the ES derived motor neurons display a significant positive correlation ($p< 0.05$) to the $Log_2$ of normalized transcript counts as determined by Spearman's rank-order correlation with a one tail distribution. Transcript counts (Ikiz et al, 2015) were normalized by gene length then to smallest value.

**Table S1:** Summary of yields and conditions for the alkylation of **3** to yield **5**.

| Batch No. | Scale | Purification Conditions | Yield (Temp) |
|---|---|---|---|
| 1 | 1.38 g, 4.30 mmol | EtOAc:Hexane (0:100 to 5:95)<br>Column length: 2.1 x 15 cm<br>Silica gel | 1.01 g, 65% (110 ºC) |
| 2 | 0.66 g, 2.05 mmol | | 0.53 g, 72% (85-90 ºC) |
| 3 | 0.43 g, 1.32 mmol | | 0.33 g, 70% (85-90 ºC) |
| 4 | 0.62 g, 1.92 mmol | | 0.49 g, 70% (85-90 ºC) |
| 5 | 0.55 g, 1.71 mmol | | 0.46 g, 75% (85-90 ºC) |

**Table S2:** Summary of yields and conditions for the deprotection of **5** to yield **6**.

| Batch No. | Scale | Purification Conditions | Yield (Temp) |
|---|---|---|---|
| 1 | 1.02g, 2.83 mmol | EtOAc:Hexane (20:80)<br>Column length: 2.1 x 10 cm<br>Silica gel | 0.43 g, 55% (50 ºC) |
| 2 | 1.0 g, 2.8 mmol | | 0.64 g, 82% (55-60 ºC) |
| 3 | 0.34 g, 0.94 mmol | | 0.22 g, 85% (60-65 ºC) |
| 4 | 0.49 g, 1.36 mmol | | 0.33 g, 89% (60-65 ºC) |
| 5 | 0.49 g, 1.36 mmol | | 0.37 g, 95% (60-65 ºC) |

**Table S3.** Summary of NMR data obtained by $^{1}H$, $^{13}C$, $^{1}H$-$^{1}H$ COSY, $^{1}H$-$^{13}C$ HMBC and 1D Selective Gradient NOESY for compound **3b**.[a]

| Position | $\delta_C$[b] type | $\delta_H$[c] ($J$ in Hz[e]) | HMBC | COSY | 1D Selective Gradient NOESY |
|---|---|---|---|---|---|
| 1 | 63.74, 63.80, $CH_2$[d] | 4.26-4.19, m<br>4.03-3.93, m<br>($^{2}J_{H,H}$ = 14.5, $^{3}J_{H,H}$ = 5.75) | 2, 3 | 2 | |
| 2 | 121.00, CH | 5.39-5.31, m ($^{3}J_{H,H}$ = 5.75) | 1, 3, 4 | 1 | |
| 3 | 139.92, 139.93, C [d] | | 1, 4, 13 | | |
| 4 | 39.31, $CH_2$ | 2.08-2.03, m | 2, 3, 5, 6, 13 | 6 | |
| 5 | 25.87, 25.91, $CH_2$ [d] | 2.21-2.11, m | 4, 6 | 6 | |
| 6 | 125.79, CH | 5.39-5.31, m<br>($^{3}J_{H,H}$ = 8.3, $^{4}J_{H,H}$ = 7.75) | 4, 5, 8, 14 | 4, 5, 14 | |
| 7 | 137.21, C | | 5, 9, 14 | | |
| 8 | 77.30, 77.32, CH [d] | 4.03-3.93, m | 6, 9, 14 | 9 | |
| 9 | 34.25, 34.27, $CH_2$ [d] | 2.31-2.22, m | 7, 8, 10, 11 | 8, 10 | |
| 10 | 120.36, CH | 5.07, t ($^{4}J_{H,H}$ = 6.5) | 8, 9, 12, 15 | 9, 12, 15 | 12 |
| 11 | 134.58, C | | 9, 10, 12, 15 | | |
| 12 | 25.99, $CH_3$ | 1.70, s | 10, 11, 12, 15 | 10 | |
| 13 | 16.47, 16.51, $CH_3$ [d] | 1.66, s | 2, 3, 4 | 2 | |
| 14 | 11.74, 11.77, $CH_3$ [d] | 1.60, s | 6, 7, 8 | 6 | |
| 15 | 18.09, $CH_3$ | 1.62, s | 10, 11, 12, 15 | 10 | |
| 1’ | 97.88, 97.99, CH [d] | 4.61, dt<br>($^{3}J_{H,H}$ = 5.0, $^{4}J_{H,H}$ = 2.8) | 1, 2’, 5’ | 5’, 3’ | |
| 2’ | 62.33, 62.35, $CH_2$ [d] | 3.49, dt<br>($^{3}J_{H,H}$ = 10.7, $^{4}J_{H,H}$ = 5.2) | 1’ 3’ | 3’, 4’ 5’ | |
| 3’ | 30.77, 30.79, $CH_2$ [d] | 1.45-1.64, m | 4’ | 1’, 2’, 4’ | |
| 4’ | 19.67, 19.68, $CH_2$ [d] | 1.45-1.64, m | 2’, 5’ | 3’, 5’ | |
| 5’ | 62.33, 62.35, $CH_2$ [d] | 3.83-3.90, m | 4’ | 3’, 4’ 2’ | |

[a]Performed in $CDCl_3$, ppm, type established by phase-sensitive HSQC. [b]125 MHz. [c]500 MHz. [d]These signals are represented as 2xs, presumably because of the two diastereotopic carbons (C8 and C1’). [e]Some $J$ values were calculated from DQF-COSY (Double Quantum Filtered COSY) experiment.

**Table S4:** Summary of yields and conditions for the oxidation of **2** to yield **3**.

| Trial Number | Scale | Purification Conditions | Yield |
|---|---|---|---|
| 1 | 4.01 g, 13.1 mmol | EtOAc:Hexane (5:95 to 20:80) | 1.4 g, 33% |
| 2 | 5.11 g, 16.6 mmol | Column length: 2.1 x 15 cm | 1.7 g, 32% |
| 3 | 4.01 g, 13.1 mmol | Silica gel | 1.38 g, 33% |
| 4 | 3.1 g, 10 mmol | | 1.2 g, 37% |

**Table S5:** Summary of yields and conditions for the bromination and subsequent diphosphorylation of **6** to yield **1**.

| Batch No. | Starting Material (Alcohol, **6**) | Cellulose Column Packing Length (cm) | Eluent | Flow Rate | Yield |
|---|---|---|---|---|---|
| 1 | 100 mg, 0.36 mmol | 2.5 x 21 | THF:0.1 M $NH_4HCO_3$ (90:10) 200 mL, then (80:20) 200 mL | 4.0-4.5 mL/min | 39 mg 25% |
| 2 | 100 mg, 0.36 mmol | 2.5 x 21 | THF:0.1 M $NH_4HCO_3$ (90:10) 200 mL, then (80:20) 200 mL | 4.0-5.0 mL/min | 42 mg 27% |
| 3 | 80 mg, 0.31 mmol | 2.5 x 21 | THF:0.1 M $NH_4HCO_3$ (90:10) 200 mL, then (80:20) 200 mL | 5.0-6.0 mL/min | 29 mg 21% |
| 4 | 100 mg, 0.36 mmol | 2.5 x 21 | THF:0.1 M $NH_4HCO_3$ (90:10) 200 mL, then (80:20) 200 mL | 5.0-6.0 mL/min | 44 mg 28% |
| 5 | 200 mg, 0.72 mmol | 2.5 x 21 | THF:0.1 M $NH_4HCO_3$ (90:10) 200 mL, then (80:20) 200 mL | 5.0-6.0 mL/min | 66 mg 21% |
| 6 | 100 mg, 0.36 mmol | 2.5 x 21 | THF:0.1 M $NH_4HCO_3$ (90:10) 200 mL, then (80:20) 200 mL | 5.0-6.0 mL/min | 42 mg 27% |
| 7 | 100 mg, 0.36 mmol | 2.5 x 21 | THF:0.1 M $NH_4HCO_3$ (90:10) 200 mL, then (80:20) 200 mL | 5.0-6.0 mL/min | 43 mg 28% |

**Table S6:** Lists of prenylated proteins enriched from ESCs, ES-MNs and ES-As.

*Included as a separate Excel File (Table S6).*

**Table S7:** Results from gprofiler enrichment analysis of prenylated proteins ESCs, ES-As and ES-MNs.

*Included as a separate Excel File (Table S7).*

### References

Ikiz B, Alvarez MJ, Ré DB, Le Verche V, Politi K, Lotti F, Phani S, Pradhan R, Yu C, Croft GF, Jacquier A, Henderson CE, Califano A, Przedborski S. The Regulatory Machinery of Neurodegeneration in In Vitro Models of Amyotrophic Lateral Sclerosis. Cell Rep. 2015 Jul 14;12(2):335-45. doi: 10.1016/j.celrep.2015.06.019. Epub 2015 Jul 2. PMID: 26146077; PMCID: PMC4646662.

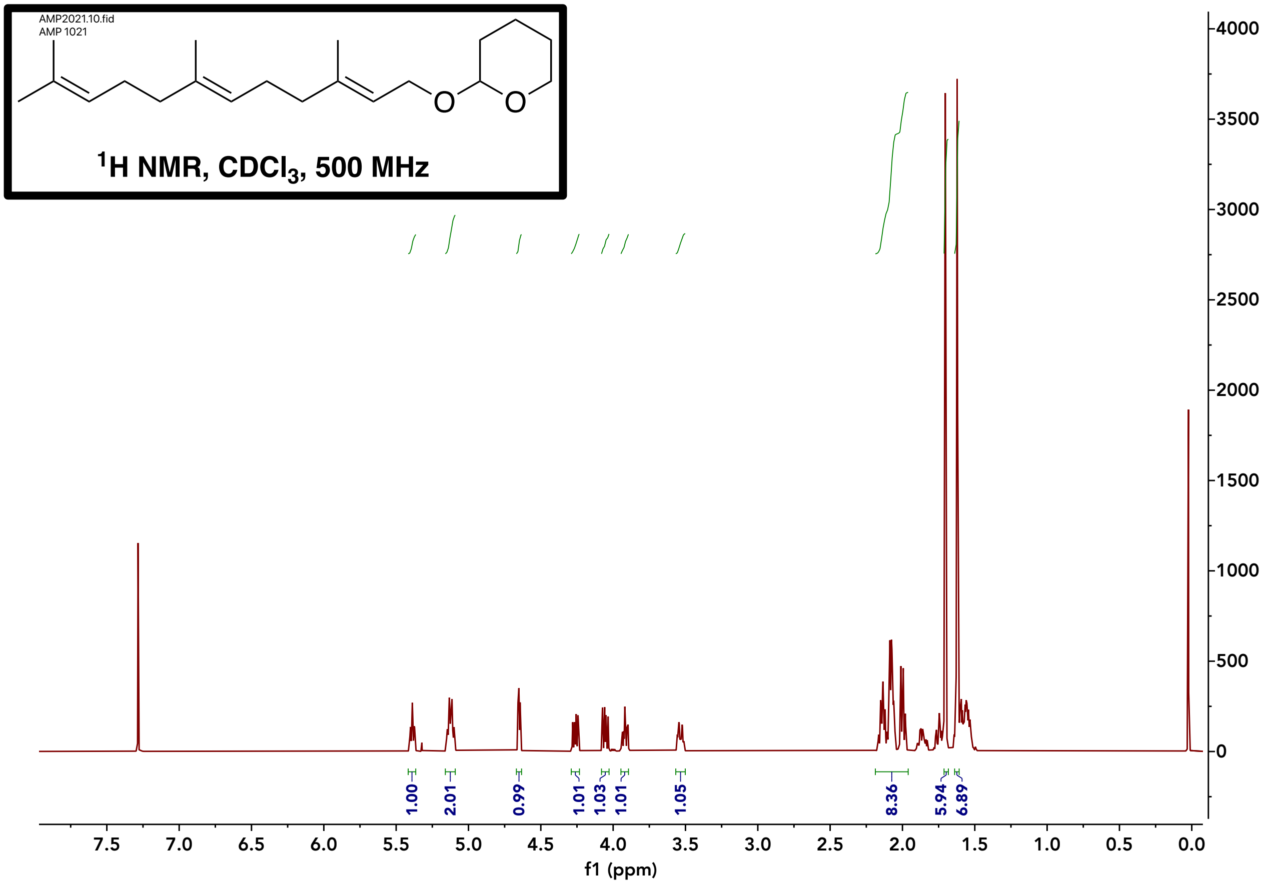

Compound **2** $^{1}$H NMR

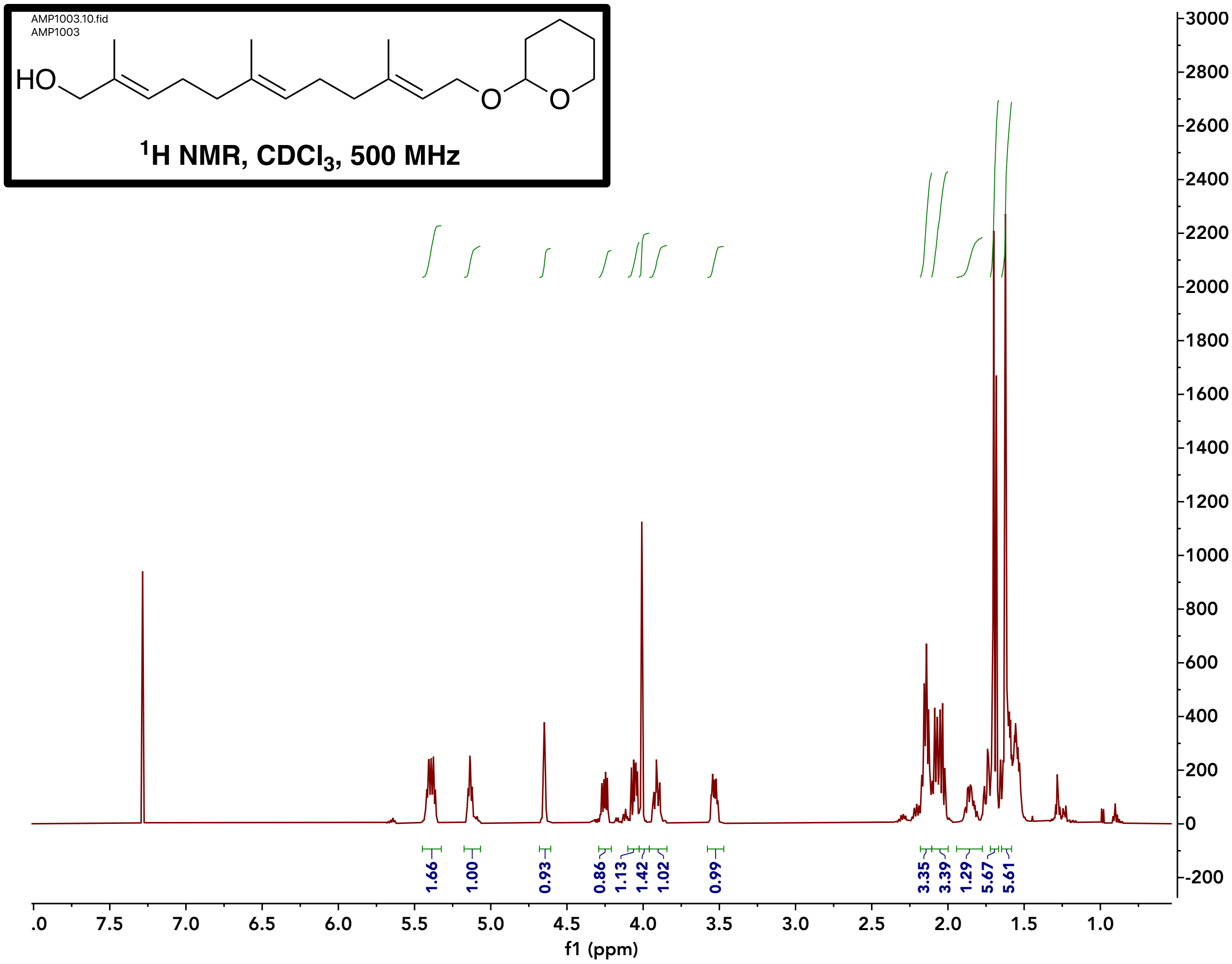

Compound **3** $^{1}$H NMR

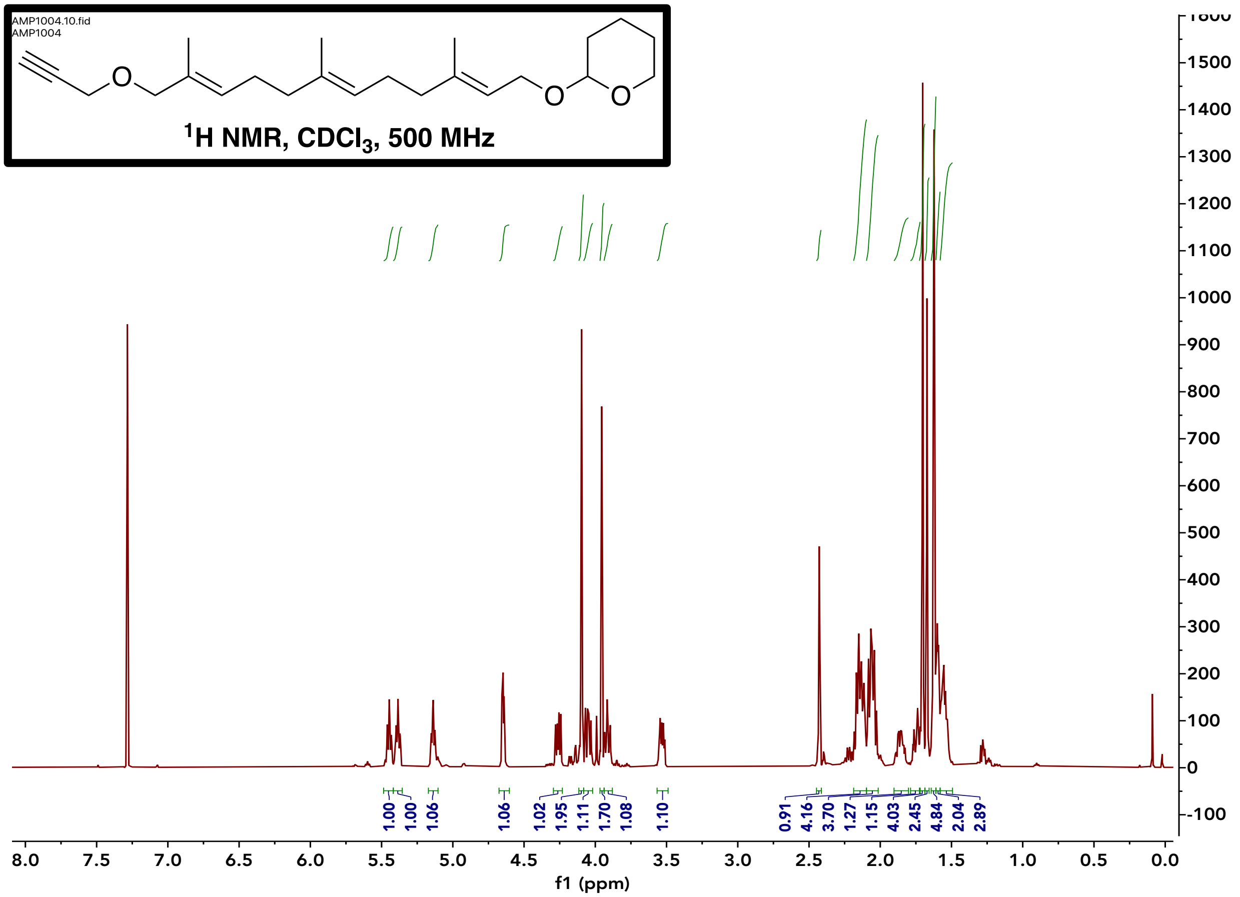

Compound **5** $^1$H NMR

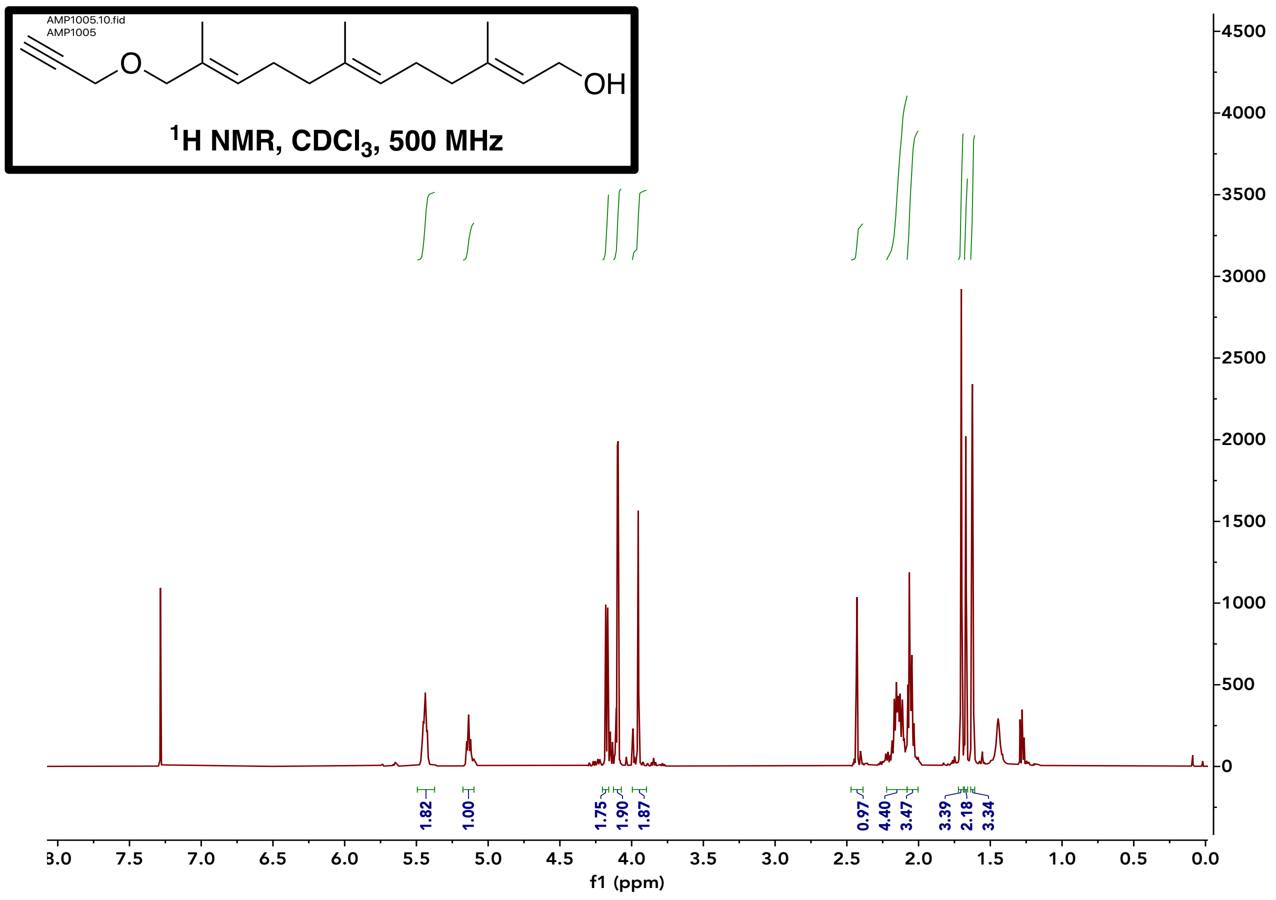

Compound **6** $^{1}$H NMR

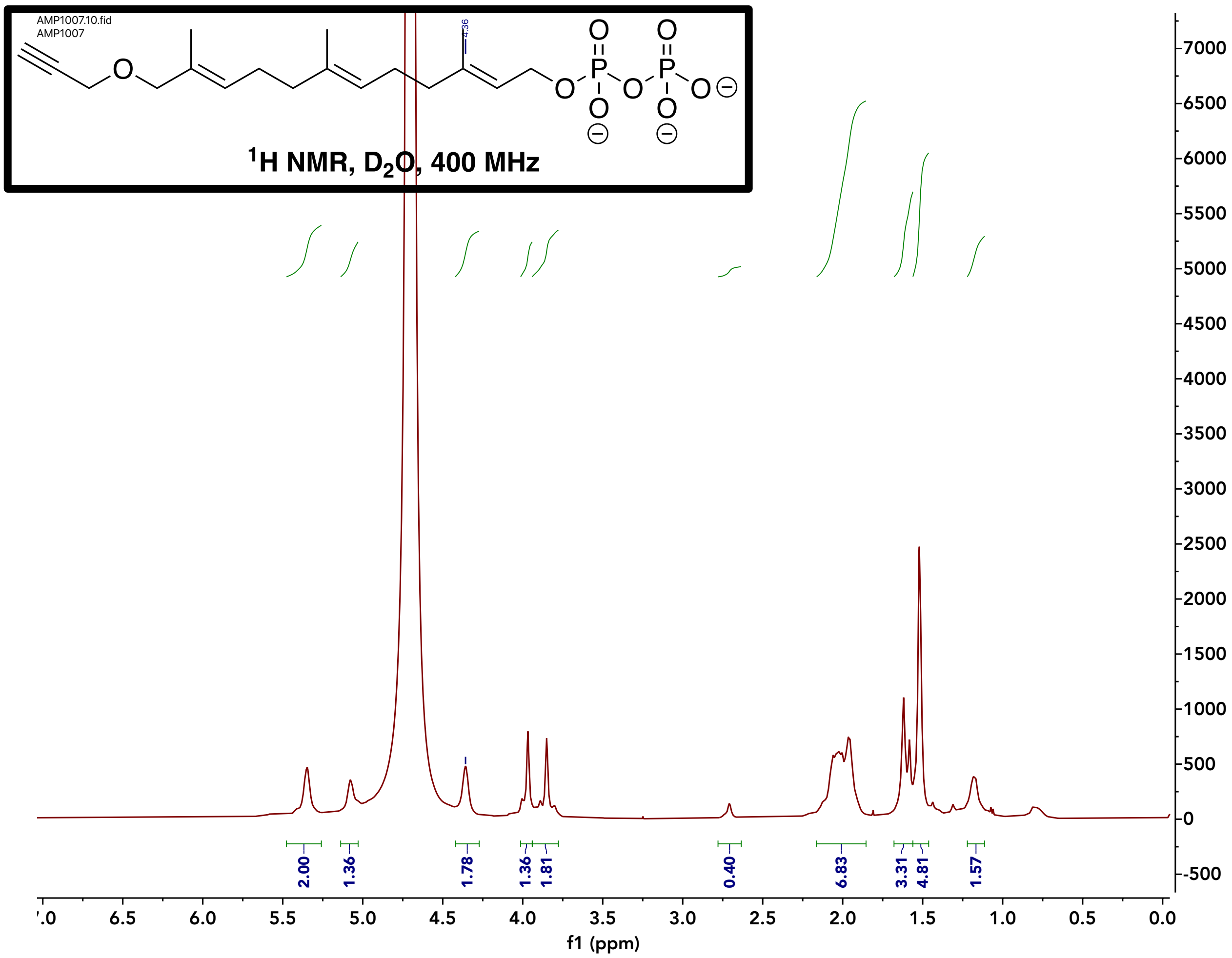

Compound **1** $^1$H NMR

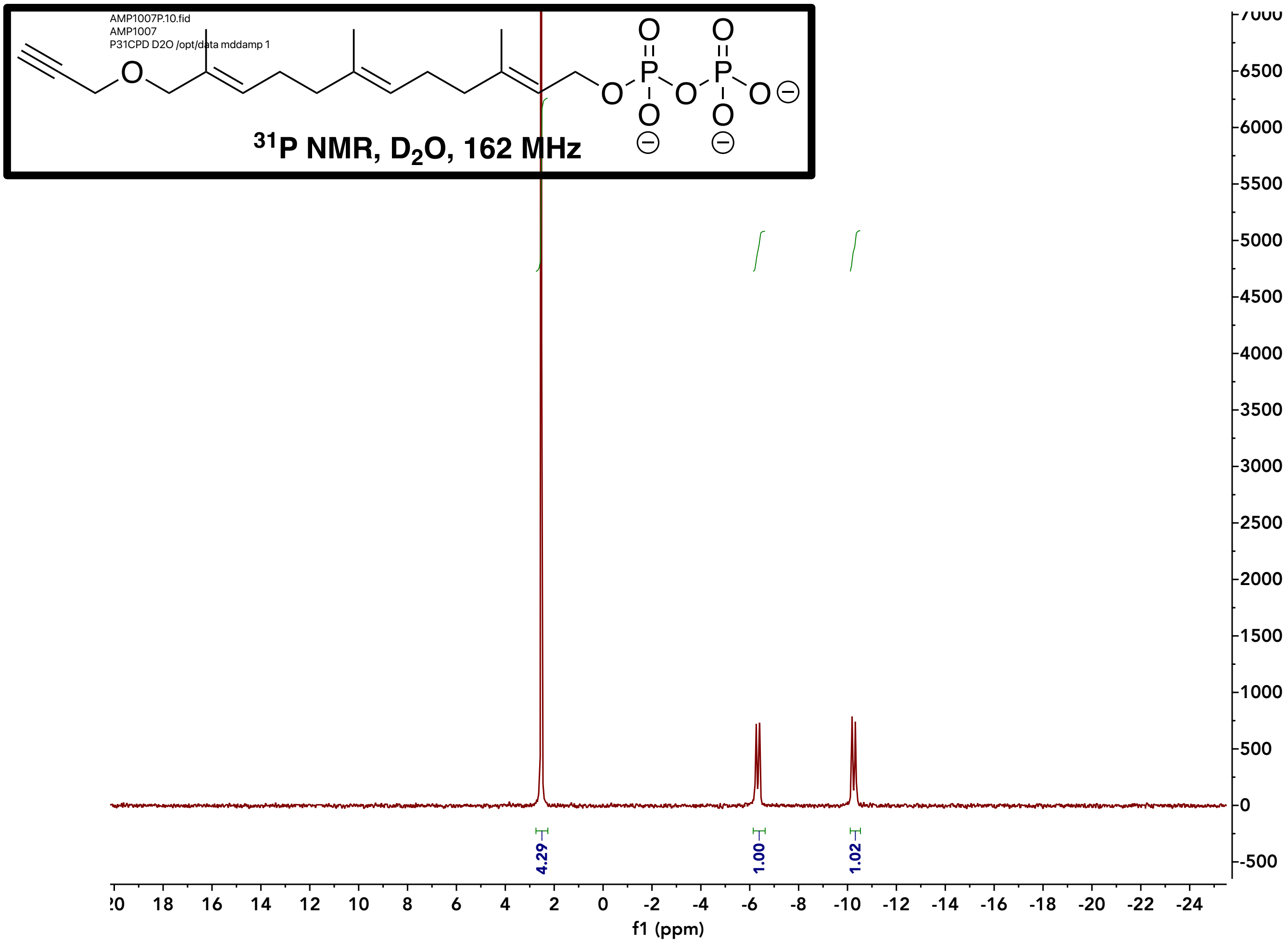

Compound **1** $^{31}$P NMR

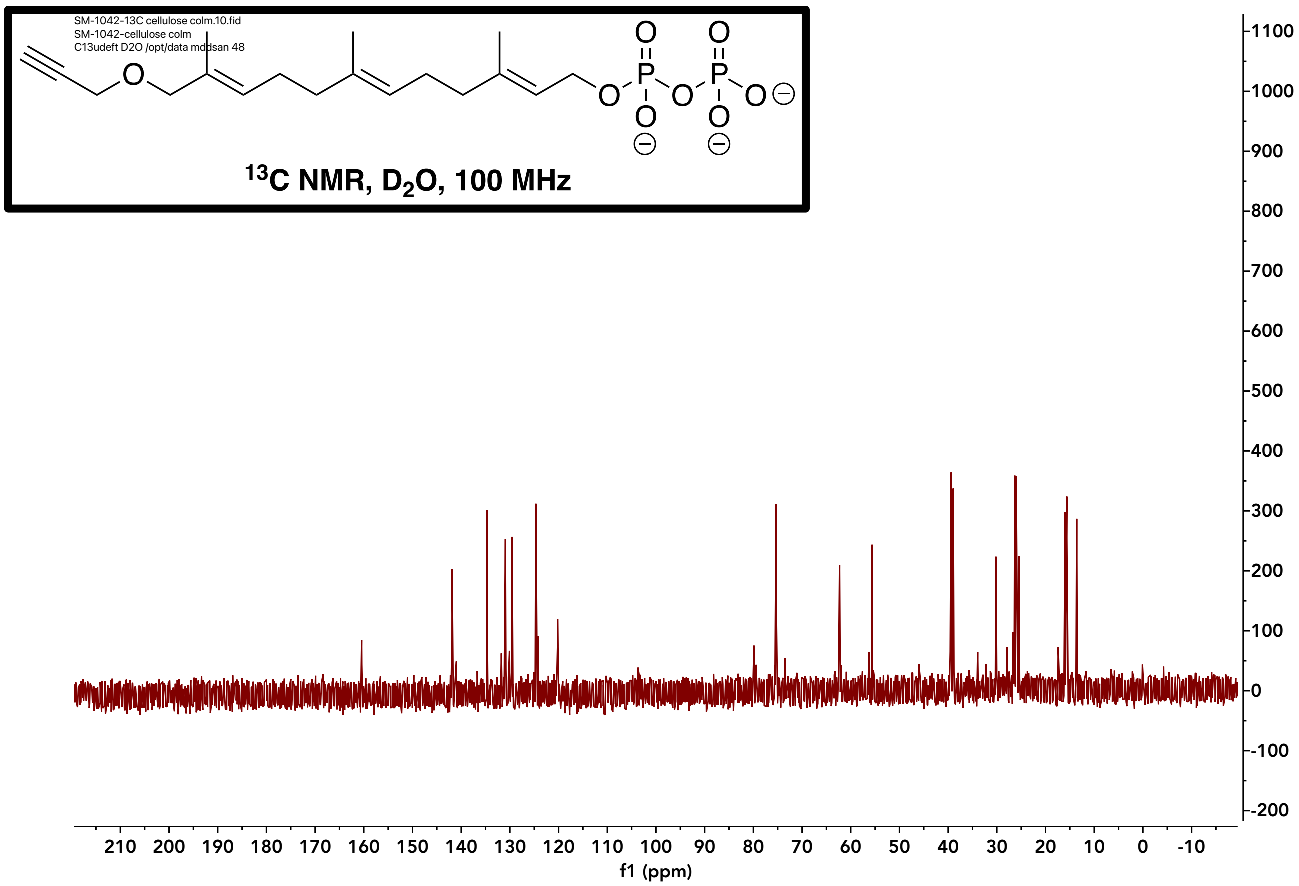

Compound **1** $^{13}$C NMR
